# Modulation of tactile feedback for the execution of dexterous movement

**DOI:** 10.1101/2021.03.04.433649

**Authors:** James M. Conner, Andrew Bohannon, Masakazu Igarashi, James Taniguchi, Nicholas Baltar, Eiman Azim

**Affiliations:** Molecular Neurobiology Laboratory, Salk Institute for Biological Studies; La Jolla, CA, USA

## Abstract

While dexterity relies on the constant transmission of sensory information, unchecked feedback can be disruptive to behavior. Yet how somatosensory feedback from the hands is regulated as it first enters the brain, and whether this modulation exerts any influence on movement, remain unclear. Leveraging molecular-genetic access in mice, we find that tactile afferents from the hand recruit neurons in the brainstem cuneate nucleus whose activity is modulated by distinct classes of local inhibitory neurons. Selective manipulation of these inhibitory circuits can suppress or enhance the transmission of tactile information, affecting behaviors that rely on movement of the hands. Investigating whether these local circuits are subject to top-down control, we identify distinct descending cortical pathways that innervate cuneate in a complementary pattern. Somatosensory cortical neurons target the core tactile region of cuneate, while a large rostral cortical population drives feed-forward inhibition of tactile transmission through an inhibitory shell. These findings identify a circuit basis for tactile feedback modulation, enabling the effective execution of dexterous movement.

## Introduction

Much of our interaction with the world occurs through movements of the hand. This “organ of considerable virtuosity” (Mountcastle 2005) achieves its impressive dexterity through dynamic interactions between motor output and sensory feedback (Johansson and Flanagan 2009, Scott 2016). Not all feedback is treated equally, however. Some sensory signals can be disruptive to behavior, for example when they are noisy, self-generated, or carry inherent temporal delays, implying circuit mechanisms for regulating the transmission of ascending information (Shadmehr, Smith et al. 2010, Scott 2016, Azim and Seki 2019). For several sensory modalities, pathways responsible for feedback modulation throughout the nervous system have begun to be defined (Sillito, Jones et al. 1994, Gilbert and Sigman 2007, Lee, Carvell et al. 2008, Fink, Croce et al. 2014, Confais, Kim et al. 2017, Liu, Latremoliere et al. 2018, Schneider, Sundararajan et al. 2018), revealing circuits that can regulate specific types of incoming signals to facilitate behavior. Studies of somatosensation have shown that injury of the pathways that carry ascending feedback from the limbs into the brain severely affects the smooth execution of dexterous behaviors (Wall 1970, Glendinning, Cooper et al. 1992, Ballermann, McKenna et al. 2001), highlighting the critical role for afferent signals from the skin and muscles in the control of movement (Johansson and Flanagan 2009). Yet, the functional organization of neural circuits that regulate the transmission of ascending somatosensory feedback and any influence this modulation has on dexterous motor output remain less clear.

The cuneate nucleus in the dorsal brainstem forms the major conduit for sensory signals from the hand ascending to the sensorimotor cortex (Berkley, Budell et al. 1986, Loutit, Vickery et al. 2020), providing a tractable location for exploring the anatomical and functional logic of feedback control. Cuneate neurons receive forelimb sensory signals directly from afferents of the dorsal root ganglia, as well as indirectly through postsynaptic dorsal column pathway neurons in the cervical spinal cord (**Fig. 1A, Suppl. Fig. 1A**) (Loutit, Vickery et al. 2020). The core, or clusters, region of the middle cuneate (hereafter referred to as Cu) mainly receives tactile input from the distal forelimbs and is thought to specialize in processing discriminative touch signals from the hand, which are likely to be involved in dexterous movements (Kuypers and Tuerk 1964, Cheema, Whitsel et al. 1983, Johansson and Flanagan 2009). Cuneolemniscal neurons in this region receive afferent input and project to several subcortical targets, most predominately the ventral posterolateral nucleus (VPL) of the thalamus, which then conveys sensory information to primary somatosensory cortex (**Fig. 1A**) (Cheema, Whitsel et al. 1983, Berkley, Budell et al. 1986). Cu has long been known to receive descending input from corticofugal projections, and cortical stimulation experiments have provided evidence for excitation and inhibition of cuneate neurons (Jabbur and Towe 1960, Andersen, Eccles et al. 1964, Kuypers and Tuerk 1964, Rustioni and Hayes 1981, Cheema, Whitsel et al. 1983, Cole and Gordon 1992, Canedo, Marino et al. 2000). Combined, these findings suggest a mechanism for top-down modulation, in which the same cortical circuits that receive forelimb sensory feedback are responsible for regulating the flow of this incoming peripheral information to the brainstem (Canedo 1997, Aguilar, Rivadulla et al. 2003), potentially through the recruitment of local inhibitory neurons (Rustioni, Schmechel et al. 1984, Popratiloff, Valtschanoff et al. 1996, Lue, Jiang-Shieh et al. 1997, Aguilar, Rivadulla et al. 2003).

**Figure 1.**
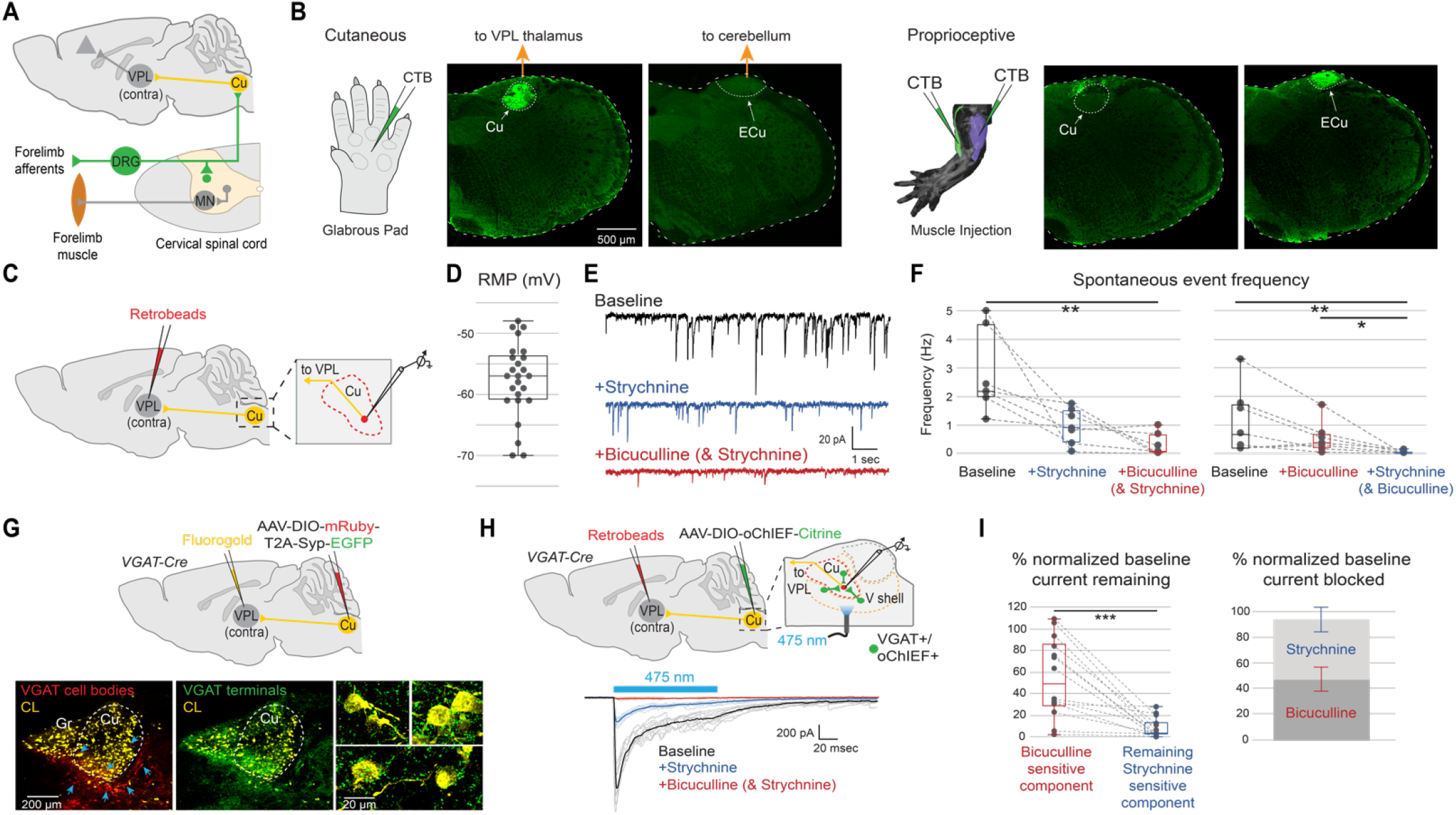
Cuneate tactile and inhibitory inputs. **(A)** Cu receives direct input from forelimb sensory afferent neurons in the dorsal root ganglia (DRG) and indirect input through ascending projections in the spinal cord (postsynaptic dorsal column pathway, not shown). Cuneolemniscal neurons project to the ventral posterolateral nucleus (VPL) of the thalamus, as well as other targets (Loutit, Vickery et al. 2020). MN, motor neuron. **(B)** Labeling direct cutaneous (left) and proprioceptive (right) projections to the dorsal column nuclei by cholera toxin B subunit (CTB) injection into peripheral end organs. Primary cutaneous afferents from the glabrous pad of the hand innervate the ipsilateral core region of the middle cuneate (Cu), but avoid the external cuneate (ECu) (6 mice). Proprioceptive afferents from forelimb muscles (biceps and triceps) rarely innervate Cu, but rather target ipsilateral ECu (right) and other regions of the cuneate nucleus (5 mice, also see **Suppl. Fig. 1B**). Major targets of Cu (VPL thalamus) and ECu (cerebellum) are indicated, though other supraspinal and spinal targets exist (Loutit, Vickery et al. 2020). Limb muscle image from (Delaurier, Burton et al. 2008). **(C)** Retrograde labeling of cuneolemniscal neurons for *in vitro* slice recording by retrobead injection into contralateral VPL thalamus. **(D)** Resting membrane potential (RMP) of labeled cuneolemniscal neurons from whole-cell recordings (25 neurons in 11 mice; current-clamp with a 0 pA holding current; all box- and whisker plots show median, 25^th^ and 75^th^ percentiles, and range). **(E)** Example traces from whole-cell recording of a cuneolemniscal neuron held at −70 mV showing spontaneous events at baseline (black). Bath application of strychnine (blue) followed by bicuculline (red) progressively eliminates spontaneous inhibitory postsynaptic currents (IPSCs). Pipette was filled with high chloride solution, causing IPSCs to appear as inward currents. **(F)** Sequential application of strychnine and bicuculline (left; 7 neurons in 4 mice; \*\**P* = 0.0015) or the reverse (right; 7 neurons in 3 mice; \*\**P* = 0.0040, \**P* = 0.0485) decreases the frequency of spontaneous IPSCs in cuneolemniscal neurons. (Friedman repeated-measures test with Dunn’s multiple comparisons test). **(G)** Viral labeling of inhibitory cell bodies (red) and their synaptic terminals (green) in *VGAT-Cre* mice and retrograde Fluorogold labeling of cuneolemniscal (CL) neurons (yellow) from VPL thalamus (2 mice). Inhibitory neurons (left; red cells, arrows) reside in the core and ventral shell regions of Cu and gracile (Gr) nucleus. Synaptic terminals (middle; green) project into the core cuneolemniscal (yellow) regions of Cu and Gr, where they make extensive contacts onto CL cell bodies (right; also see **Suppl. Fig. 3**). **(H)** Slice recording from CL neurons retrogradely labeled by retrobead injection into contralateral VPL thalamus (top). Cu inhibitory neurons were optogenetically activated following viral expression of oChIEF in Cu core and ventral shell (V shell) regions of *VGAT-Cre* mice. Example whole-cell recording from a labeled CL neuron (bottom). Photoactivation of Cu inhibitory neurons elicits large IPSCs that are eliminated by sequential application of strychnine and bicuculline. Faint lines represent single trials, dark lines represent mean. **(I)** Sequential application of bicuculline and strychnine shows cumulative reduction in the amplitude of light-evoked IPSCs (left, normalized to baseline) with an approximately equal mix of GABA and glycine mediated components (right, error bars indicate SEM). (14 neurons in 7 mice; \*\*\**P* = 0.0001; Wilcoxon two-tailed matched-pairs signed rank test; also see **Suppl. Fig. 5C**).

However, lack of circuit specific access has limited efforts to define the organization of these inhibitory neurons, the relationship they might have to descending cortical pathways, and any impact their putative feedback modulation might have on dexterous forelimb movement. Stated more generally, it remains unclear which neural circuits regulate the incessant flow of sensory signals to ensure that only the appropriate and salient information is used to impact ongoing behavior. Here, with a focus on tactile signaling in the Cu of mice, we sought to define the organization, functional connectivity, and behavioral implications of circuits that regulate feedback from the hand as it enters the brain.

## Results

### Cuneate tactile and inhibitory inputs

We first localized the region of the middle cuneate that receives direct tactile input from the hand for subsequent anatomical, electrophysiological, and behavioral experiments. Cutaneous afferents were targeted at their peripheral terminals by injection of cholera toxin B subunit (CTB) into the glabrous pad of the hand, revealing dense innervation of the ipsilateral Cu, which sends its most prominent output to contralateral VPL thalamus (**Fig. 1B**) (Berkley, Budell et al. 1986). Conversely, proprioceptive afferents were targeted by injection of CTB into forelimb muscles, revealing minimal innervation of Cu, but some targeting of other cuneate regions and dense projections to the neighboring ipsilateral external cuneate nucleus (ECu), which mainly projects to the cerebellum (**Fig. 1B**) (Loutit, Vickery et al. 2020). We confirmed these findings by genetically restricting our tracing to proprioceptors through conditional viral targeting of forelimb muscles in *Pv-Cre* mice (**Suppl. Fig. 1B**). Supporting previous work (Hantman and Jessell 2010, Niu, Ding et al. 2013), these results establish that direct ascending cutaneous and proprioceptive afferents remain largely segregated at the level of the dorsal column nuclei in mice.

Several studies have shown that sensory responses in Cu are subject to attenuation, potentially mediated by local inhibition (Andersen, Eccles et al. 1962, Andersen, Eccles et al. 1964, Rustioni, Schmechel et al. 1984, Lue, Jiang-Shieh et al. 1997, Aguilar, Rivadulla et al. 2003). Yet the detailed organization of local circuits that can affect tactile transmission, and any contribution they might have to movement are not well understood. Having localized the tactile recipient Cu region, we next sought to characterize the properties of cuneolemniscal neurons in Cu by performing whole-cell electrophysiological recordings from adult brainstem slice preparations (**Fig. 1C**). We found that retrogradely labeled cuneolemniscal neurons have a relatively depolarized resting membrane potential (mean = −57.60 mV ± 1.22 mV SEM; **Fig. 1D**) that is on average slightly below the action potential threshold (mean = −49.67 mV ± 1.78 mV SEM; **Suppl. Fig. 2A**), similar to the high resting membrane potential found in cat cuneate neurons (Bengtsson, Brasselet et al. 2013). Moreover, we found that these neurons receive extensive spontaneous inhibitory input that is abolished by the combined application of the glycine receptor antagonist strychnine and the GABA_A_ receptor antagonist bicuculline (**Fig. 1E,F** and **Suppl. Fig. 2B**). These results indicate that Cu neurons that convey tactile information to the thalamus are broadly inhibited by GABAergic and glycinergic inputs.

To identify the location of neurons that might provide this inhibition, we genetically restricted fluorophore expression to GABAergic and glycinergic neurons through conditional viral targeting of the Cu region of *VGAT-Cre* mice. We found inhibitory neurons within the core region of the Cu, where cuneolemniscal neurons reside, as well as throughout a shell region ventral to the Cu core, where cuneolemniscal neurons are absent (collectively referred to as Cu inhibitory neurons, **Fig. 1G**). Visualization of a synaptically tagged fluorophore revealed that axon terminals arising from these local inhibitory neurons project heavily into the Cu core region, where they provide dense synaptic input onto cuneolemniscal neurons (**Fig. 1G**). Genetic labeling of inhibitory subclasses revealed that glycinergic neurons are present throughout the Cu core and shell regions, whereas GABAergic neurons reside mostly in the ventral shell (**Suppl. Fig. 3A,B**). Monosynaptic retrograde rabies tracing originating specifically from Cu neurons that target VPL thalamus confirmed that both GABAergic and glycinergic neurons directly innervate cuneolemniscal neurons (**Suppl. Fig. 3C,D**). Together, these findings identify the localization and connectivity of inhibitory neurons in the cuneate, leading us to ask whether they do in fact modulate cuneolemniscal activity.

To assess the functional connectivity of these circuits, we expressed the excitatory opsin oChIEF in Cu inhibitory neurons through conditional viral injection in *VGAT-Cre* mice and performed whole-cell electrophysiological recording from adult brainstem slices. Confirming the efficacy of this optogenetic approach, targeted inhibitory neurons showed robust responses to photoactivation (**Suppl. Fig. 4A-C**). Moreover, these inhibitory neurons receive extensive spontaneous inhibitory inputs that appear to be largely GABAergic (**Suppl. Fig. 4D-F**), supporting previous immunohistochemical evidence of local inhibitory connectivity in rats (Lue, Jiang-Shieh et al. 1997), and suggesting a means for local disinhibition. We next recorded postsynaptic responses from tagged cuneolemniscal neurons and found that optogenetic activation of local Cu inhibitory neurons produces large inhibitory currents (**Fig. 1H**) with onset kinetics indicative of monosynaptic connectivity (**Suppl. Fig. 5A,B**). Combined application of strychnine and bicuculline abolished light-evoked responses (**Fig. 1H,I**), with each cuneolemniscal neuron showing a distinct mix of GABAergic and glycinergic inputs (**Fig. 1I** and **Suppl. Fig. 5C**). A previous model proposed that GABAergic cells inhibit cuneolemniscal neurons while glycinergic neurons disinhibit cuneolemniscal neurons by inhibiting GABAergic cells (Aguilar, Rivadulla et al. 2003, Soto, Aguilar et al. 2004). Our results expand upon and revise this model by demonstrating that both GABAergic and glycinergic cells elicit direct cuneolemniscal inhibition in varying combinations.

### Local inhibitory modulation of ascending tactile feedback

While these findings establish the anatomical and functional relationship between Cu inhibitory neurons and cuneolemniscal neurons, they do not demonstrate whether these circuits can modulate tactile signaling. To address this question, we performed *in vivo* extracellular electrophysiological recordings from tactile-responsive neurons in the Cu of anesthetized mice while physical stimuli were applied to the glabrous pad of the ipsilateral hand by a rotating wheel (**Fig. 2A**). Measurement of evoked activity within Cu allowed us to identify neurons that are robustly sensitive to tactile input (**Fig. 2B,C**).

**Figure 2.**
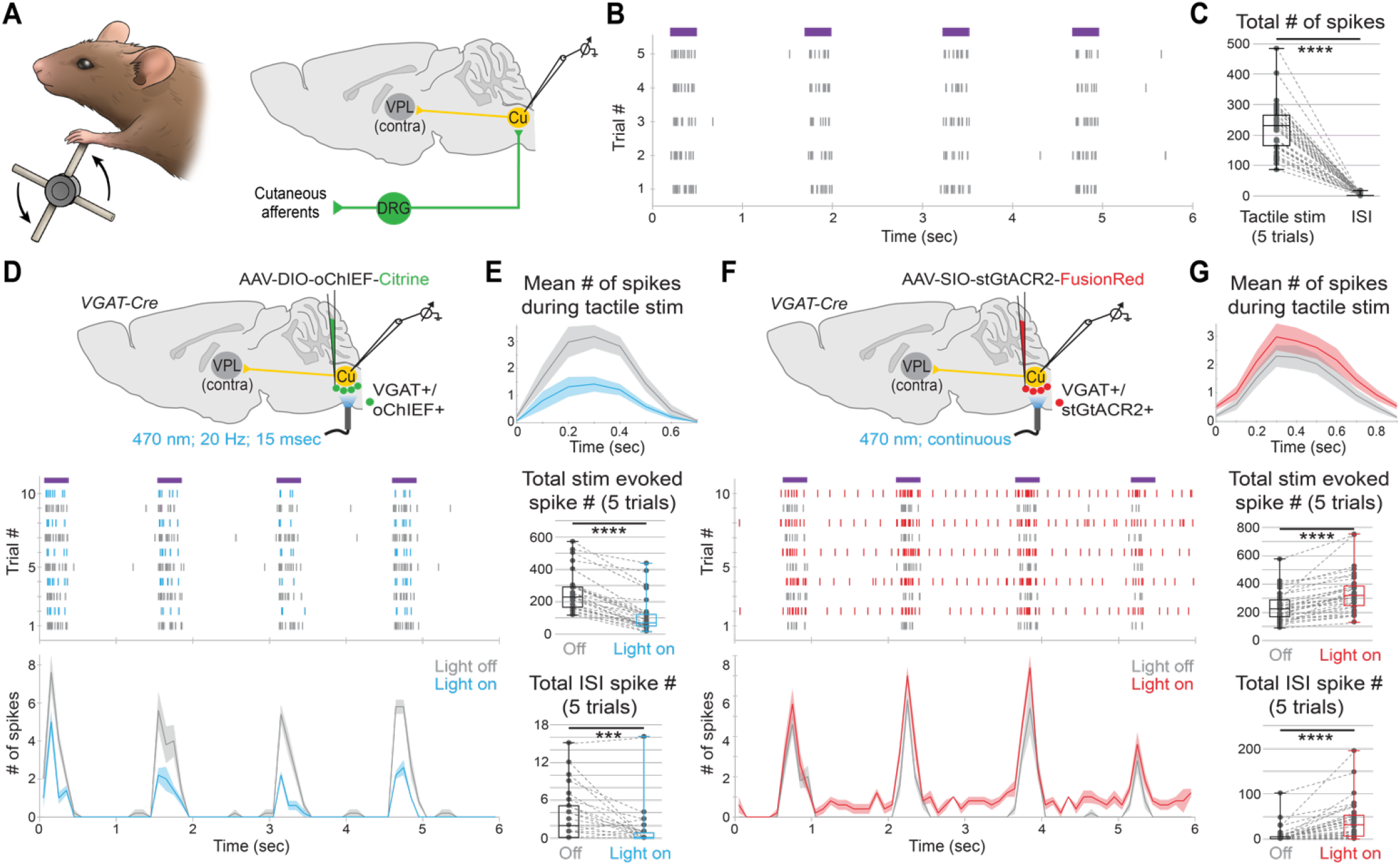
Local inhibitory circuits bidirectionally modulate tactile responses. **(A)** *In vivo* extracellular recording in Cu of anesthetized mice with tactile stimuli applied to the ipsilateral glabrous pad of the hand (see Materials and Methods). **(B)** Spike raster plots of tactile-evoked signals from an example Cu recording site across five trials with four stimuli per trial (20 total stimuli). Purple bars indicate periods when tactile stimulus is applied. **(C)** Total spikes across all 20 stimuli for each recording site show robust tactile-evoked responses with little spontaneous activity during the inter-stimulus intervals (ISI). (51 recordings across 3 mice; \*\*\*\**P* < 0.0001; Wilcoxon two-tailed matched-pairs signed rank test). **(D)** *In vivo* extracellular recording of tactile responsive units in Cu in anesthetized mice (top). Local Cu inhibitory neurons were targeted for optogenetic activation through viral oChIEF expression in *VGAT-Cre* mice. Spike raster plots (middle) and mean spike number histograms (bottom; 0.1 sec bin) from an example recording site across ten interleaved trials (five light off, gray; five light on, blue; each with four tactile stimuli, purple bars), show a suppression of tactile-evoked spikes during photoactivation of Cu inhibitory neurons. Lines in lower histogram indicate mean, shaded areas represent SEM. **(E)** Average mean number of spikes during a single tactile stimulus (top; 0.1 sec bin) across all recordings (30 sites from 3 mice) shows suppression of tactile-evoked responses in Cu neurons during photoactivation of local inhibitory neurons. Lines indicate mean, shaded areas represent SEM. Pairwise comparisons of the total number of spikes across five trials (20 tactile stimuli each for light off and light on) reveal suppression of tactile-evoked spikes (middle; \*\*\*\**P* < 0.0001) and suppression of spontaneous activity during tactile ISI periods (bottom; \*\*\**P* = 0.0004). Note the different scales for tactile-evoked and ISI activity (Wilcoxon two-tailed matched-pairs signed rank test). **(F)** *In vivo* extracellular recording (top; as in (D)). Local Cu inhibitory neurons were targeted for optogenetic inhibition through viral stGtACR2 expression in *VGAT-Cre* mice. Spike raster plots from an example recording site (middle; as in (D); light off, gray; light on, red) show an increase in spikes during photoinhibition of Cu inhibitory neurons. Spike increase is especially apparent during the tactile ISI periods. Lines in histogram (bottom) indicate mean, shaded areas represent SEM. **(G)** Quantification (as in (E)) across all recordings (34 sites from 3 mice) shows an increase in tactile-evoked responses in Cu neurons during photoinhibition of local inhibitory neurons (top). Pairwise comparisons reveal an increase in spikes evoked by tactile stimuli (middle; \*\*\*\**P* < 0.0001) and an increase in spontaneous activity during tactile ISI periods (bottom; \*\*\*\**P* < 0.0001). (Wilcoxon two-tailed matched-pairs signed rank test).

We then asked how the transmission of tactile signals from the ipsilateral hand is affected by activation or inactivation of local Cu inhibitory circuits. First, conditional viral expression of oChIEF in *VGAT-Cre* mice was used to assess the impact of activating Cu inhibitory neurons on tactile-responsive neurons (**Fig. 2D**). Pairwise comparisons show that during photostimulation, the number of tactile-evoked spikes was consistently suppressed (60.99% ± 3.71% SEM reduction in spike number), and this attenuation applied even to the rare spontaneous activity observed during tactile inter-stimulus periods (56.93% ± 15.40% SEM reduction in spike number) (**Fig. 2D,E**). Next, we used conditional viral expression of the inhibitory opsin stGtACR2 in *VGAT-Cre* mice to evaluate the impact of inactivating Cu inhibitory neurons (**Fig. 2F**). We found that photoinhibition of Cu inhibitory neurons amplified the number of tactile-evoked spikes (50.46% ± 8.87% SEM increase in spike number), and elicited a striking increase in inter-stimulus spiking activity (1,333.74% ± 354.81% SEM increase in spike number), suggesting that local inhibition prevents aberrant Cu neuronal firing and the transmission of spurious sensory information (**Fig. 2F,G**). Together, these findings indicate that recruitment or suppression of Cu inhibitory neurons provides a circuit basis for bidirectional modulation of tactile signaling through the cuneate.

### Disrupting cuneate modulation perturbs dexterous movements

The regulation of feedback is a fundamental aspect of coordinated behavior (Scott 2016, Azim and Seki 2019), and dexterous movements might be particularly dependent on the dynamic adjustment of sensory signaling (Johansson and Flanagan 2009). Dorsal column lesion studies suggest that fine movements are especially susceptible to the loss of somatosensory information (Wall 1970, Glendinning, Cooper et al. 1992, Ballermann, McKenna et al. 2001, Mountcastle 2005), but lack of temporal and circuit specificity in these experiments leave any role for feedback modulation unclear. The genetic access to Cu inhibitory circuits that we established provided us the opportunity to answer a question that classical lesion studies cannot address – does the regulation of tactile feedback as it ascends into the brain have any impact on dexterous motor control?

As a first assay, mice were trained to perform a string pulling behavior that elicits the smooth alternation of left and right hands as the animal reaches, grasps, and pulls, mimicking many natural behaviors (Blackwell, Banovetz et al. 2018) (**Fig. 3A**). We reasoned that this assay could help to distinguish two possible scenarios: spinal tactile reflex circuits are sufficient for the smooth execution of string grasping during this rhythmic behavior, or alternatively, ascending tactile signals need to be regulated appropriately in the cuneate for effective performance. To attenuate tactile signaling in the cuneate, we targeted viral delivery of oChIEF to Cu inhibitory neurons of *VGAT-Cre* mice (**Fig. 3A**) and found that photoactivation of these neurons affected the animals’ ability to coordinate ipsilateral grasping movements with string contact (**Fig. 3B,C**). Automated tracking revealed that these prehension mistakes affected limb kinematics, causing a greater number of hand direction reversals after grasping errors and a reduction in movement path length as the animals made successive attempts to correctly time the grasp (**Fig. 3D**, **Supplementary Movies 1,2**). Prehension errors and kinematic deficits did not appear in the contralateral limb (**Fig. 3B-D**) nor in control mice receiving photostimulation (**Suppl. Fig. 6**). Conversely, to evaluate how an aberrant increase in tactile signaling affects behavior, we expressed the inhibitory opsin stGtACR2 in Cu inhibitory neurons of *VGAT-Cre* mice. During photoinhibition, approximately half of the mice appeared to execute the behavior normally, while the other half often halted string pulling movements, and in the more extreme cases, dropped the string and gripped the ipsilateral hand (**Supplementary Movie 3**). These findings suggest that eliciting aberrant tactile transmission through the cuneate can disrupt the smooth execution of movement, though this behavior might be somewhat resilient to spurious sensory signals, perhaps due to the reward-driven nature of the task. More generally, these combined results indicate that spinal tactile circuits alone are not sufficient for coordinating grasping, and a shift in the balance of ascending tactile transmission in either direction can severely impact the execution of sensory-guided behaviors.

**Figure 3.**
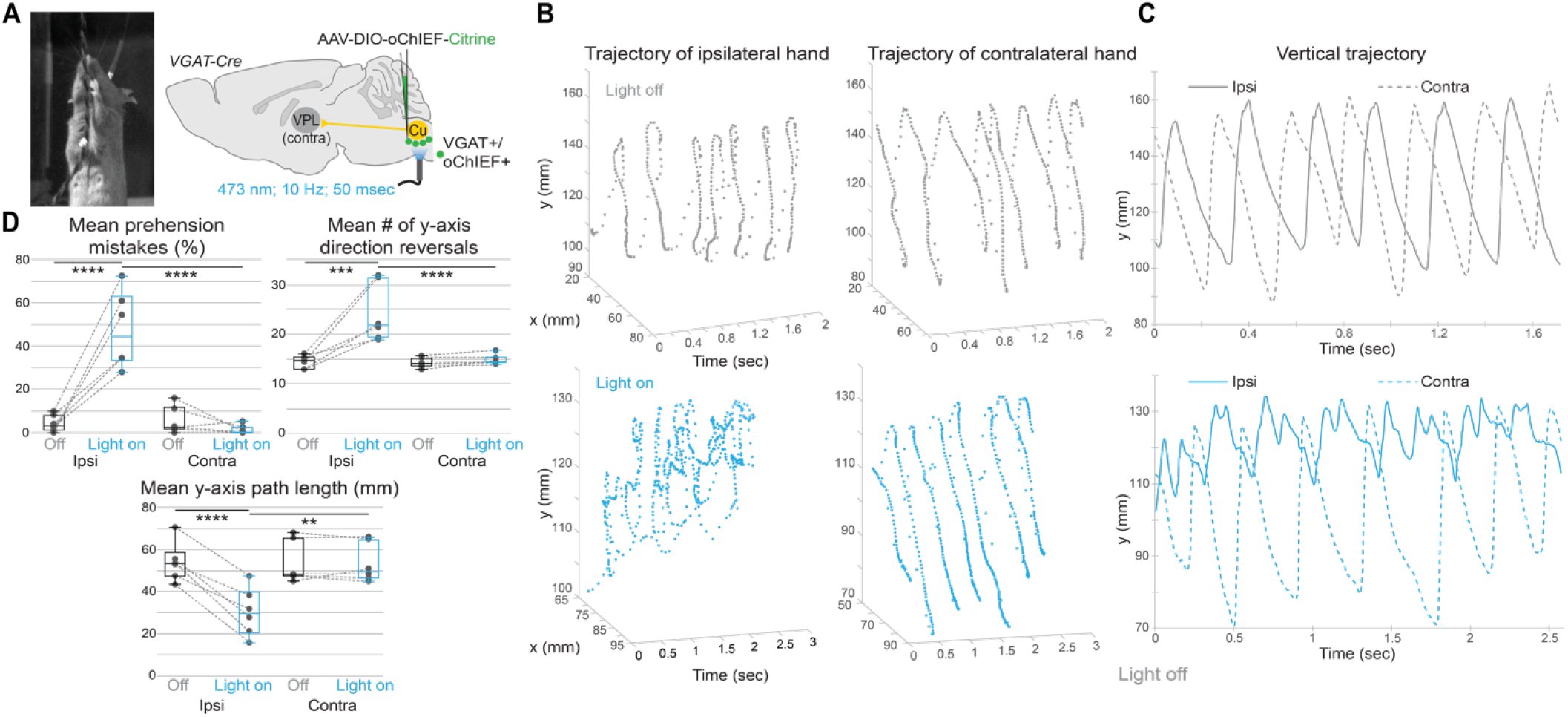
Perturbing cuneate inhibitory circuits disrupts dexterous movements. **(A)** String pulling task (left). Local Cu inhibitory neurons were targeted for optogenetic activation through viral oChIEF expression in *VGAT-Cre* mice. **(B)** Example trajectories of ipsilateral (left) and contralateral (right) hands in horizontal (x) and vertical (y) dimensions over time. With the light off (gray), both hands exhibit smooth cycles with uninterrupted pulling paths. During photoactivation (blue), ipsilateral kinematics exhibit frequent interruptions in the pulling paths and decreased pulling distance per cycle, while the contralateral limb is unaffected. **(C)** Example y trajectories of ipsilateral (solid) and contralateral (dashed) hands. With the light off (gray), both hands exhibit a smooth, alternating trajectory. During photoactivation (blue), the contralateral limb is unaffected, while the ipsilateral hand exhibits frequent direction reversals with short pull distances at the top of the trajectory, reflecting corrective attempts after prehension errors. **(D)** Quantification of prehension errors and kinematics across 6 mice. Prehension errors of the ipsilateral, but not contralateral, hand increase during photoactivation (top left; % of grasp attempts in which an error was made across trials, see Materials and Methods), as do the mean number of ipsilateral vertical (y) direction reversals (top right; mean number of direction reversals per trial). The mean y path length traversed by the ipsilateral hand between direction reversals decreases during photoactivation (bottom; mean absolute distance between the peak and trough of a given path segment across trials) (\*\*\*\**P* < 0.0001, \*\*\**P* = 0.0003, \*\**P* = 0.0016; Two-way repeated measures ANOVA with Sidak multiple comparisons test).

While the string pulling assay provides an ethologically relevant means to evaluate goal-directed limb movements, animals in this task are free to use any combination of tactile, proprioceptive, and visual feedback. We next wanted to probe the tactile component of a dexterous movement more selectively. Human studies have shown that tactile acuity is surprisingly high during object manipulation when compared to psychometric detection thresholds, suggesting that feedback used in the service of movement need not rise to the level of perceptual awareness (Pruszynski, Flanagan et al. 2018). Thus, we reasoned that a fine motor control assay could provide the most sensitive way to evaluate the impact of Cu feedback modulation. Inspired by these human behavioral experiments (Pruszynski, Flanagan et al. 2018), we developed a novel quantitative tactile orienting assay for mice that enabled us to isolate the contribution of salient cutaneous feedback to task execution. Head-fixed mice were trained to use their hand to turn a pedestal with an oriented texture cue to a defined target zone and hold in place (**Fig. 4A-D**, see Materials and Methods). The pedestal was affixed to a motor/rotary encoder assembly, enabling the pedestal texture to be set to a random starting orientation at the beginning of each trial (**Suppl. Fig. 7A**). Over the course of 4-6 weeks of training, the task conditions (range of starting orientation, target zone, hold duration) gradually became more difficult in a closed-loop fashion, depending on each animal’s behavioral performance, until expert performance was achieved at the most stringent conditions (defined as at least 60% success, usually exceeding 90%, see Materials and Methods; **Fig. 4E**).

**Figure 4.**
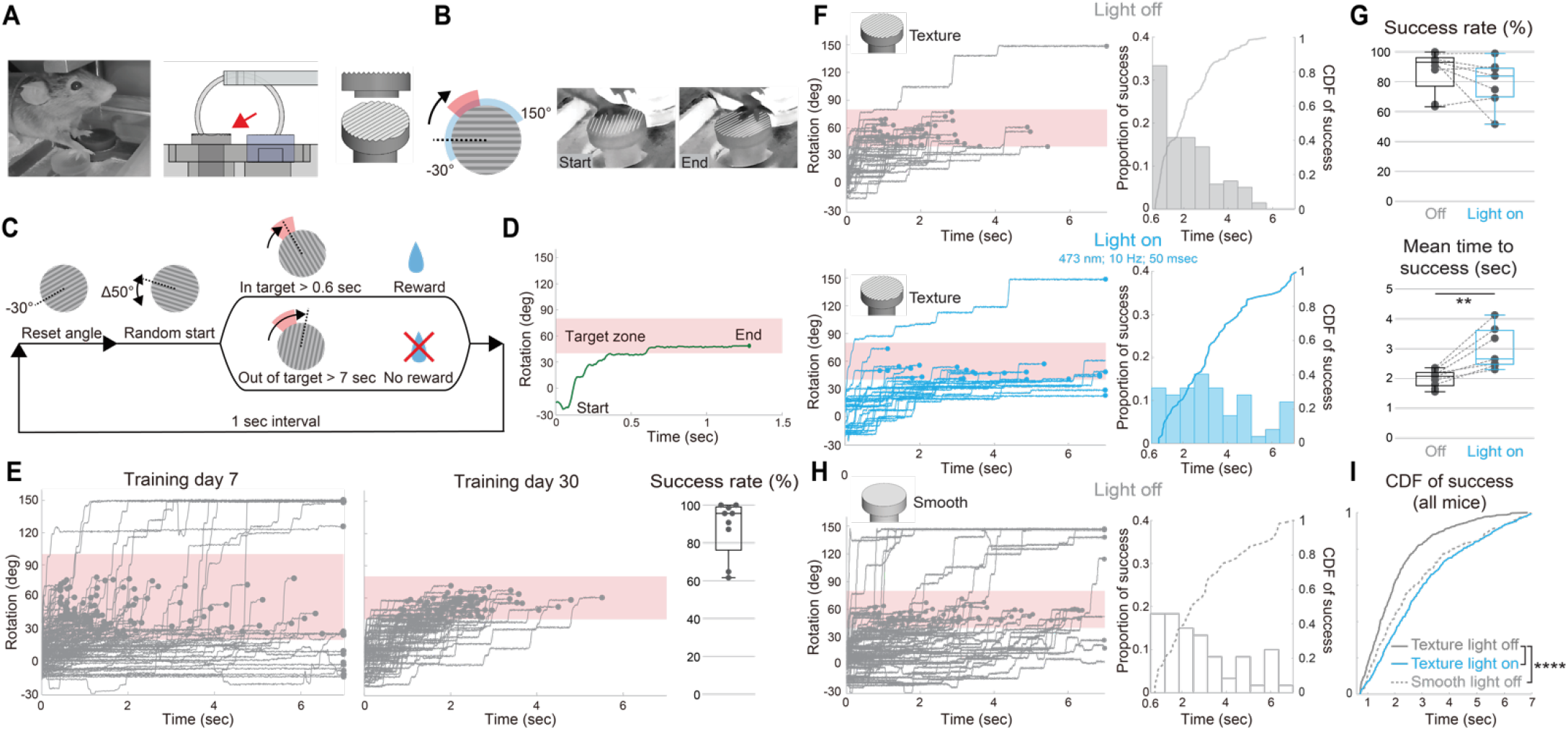
Activation of cuneate inhibitory circuits compromises tactile-dependent movements. **(A)** Head-fixed mouse performing the tactile orienting task (left, see Materials and Methods). A pedestal (middle, red arrow) with parallel orientation ridges (right) was placed under the right hand. The pedestal was connected to a motor/rotary encoder assembly (also see **Suppl. Fig. 7A**). The left hand was placed on a fixed surface of equal height. **(B)** The neutral angle of the ridges was defined as perpendicular to the body axis of the mouse (0°, dotted line), and the pedestal was passively mobile through a 180° range (blue, −30° to 150°). To receive water reward, the animal must turn the pedestal clockwise to align the ridges within a defined target range (red). **(C)** Task structure. After training, at the beginning of each trial, the pedestal is reset to −30°, and then adjusted to a random starting orientation (0° ± 25°). The pedestal then becomes passively mobile, indicating the start of the trial. Water reward is delivered if the orientation of the ridges stays within the target zone (60° ± 20°) for > 0.6 sec. Otherwise, the trial is terminated without reward after 7 sec. The next trial begins after a 1 sec interval, and trials continue until either 105 rewards are delivered or 160 trials are completed. **(D)** Example of angular trajectory of pedestal during a single trial. **(E)** Representative angular trajectories during training. Animals begin training with a small starting range and large target range (left). As training proceeds, the starting range gets larger and target zone gets smaller (middle; see Materials and Methods). Mean success rate after training (right, 9 mice). **(F)** Ipsilateral Cu inhibitory neurons were targeted for optogenetic activation (as in Fig. 3A). Example angular trajectories (left) with light off (top; gray) and light on (bottom; blue), and plots (right) showing the proportion of successful trials at each time point (bars, 0.64 sec bins) and the cumulative distribution function (CDF) for only the successful trials (lines). **(G)** During optogenetic activation (blue), the overall success rate is unaffected (top), but the mean elapsed time to achieve success increases (bottom; 7 mice; \*\**P* = 0.0047; Two-way mixed-effects model with Geisser-Greenhouse correction and Sidak multiple comparisons test; also see **Suppl. Fig. 7B,C**). **(H)** Example angular trajectories, binned successes, and CDF (as in (F)), for a mouse performing the task with a smooth surface (no ridges) and the light off. **(I)** CDF across all three conditions shows an equivalent drop in performance from control conditions (texture with light off, gray line, 9 mice) when photoactivating Cu inhibitory neurons (blue line, 7 mice) or when removing texture (dashed gray line, 8 mice) (\*\*\*\**P* < 0.0001; Kruskal-Wallis test with Dunn’s multiple comparisons test; also see **Suppl. Fig. 7B,C** and Materials and Methods).

To determine how disrupting the modulation of tactile feedback in the cuneate affects task performance, we expressed oChIEF in Cu inhibitory neurons (**Suppl. Fig. 7B**). Photoactivation did not affect overall task success in trained mice (when given a 7 sec time window, see Materials and Methods), but did result in an increase in the amount of time taken to reach the target, appearing as a rightward shift in the cumulative distribution of successes over time (**Fig. 4F,G,I** and **Suppl. Fig. 7B,C**). To determine how these deficits compare to normal performance when tactile information is more impoverished, we evaluated task performance using a smooth pedestal lacking tactile ridges. Unperturbed animals using a smooth platform also showed a rightward shift in the cumulative distribution of successes over time (**Fig. 4H** and **Suppl. Fig. 7B,C**). These findings suggest that this performance gap represents the sensory advantage provided by salient tactile cues, and in their absence, mice rely on other sensory modalities or more exploratory behavioral strategies to find the target. Most notably, animals orienting a smooth pedestal showed a nearly identical behavioral deficit to those using a textured platform during Cu inhibitory neuron activation (**Fig. 4I**). Behavioral performance was not affected in control animals receiving photostimulation (**Suppl. Fig. 7D**). Together, our behavioral findings confirm the essential role for ascending tactile signals in dexterous behavior, and reveal that dysfunction of Cu modulatory circuits impairs effective interaction with the environment.

### Top-down corticofugal control of cuneate circuits

In a final set of experiments, we asked whether the modulatory circuits we identify can provide a circuit basis for top-down control of tactile feedback. Ascending sensory transmission is attenuated before and during limb movement (Ghez and Pisa 1972, Coulter 1974, Chapin and Woodward 1981), potentially as a means to gate reafferent feedback caused by one’s own movement or increase the signal-to-noise of sensory information most relevant to task execution (Chapman 1994, Azim and Seki 2019). The primary somatosensory cortex (SSp) has long been known to innervate the cuneate nucleus, at least in part through collaterals of corticospinal projection neurons (Kuypers and Tuerk 1964, Rustioni and Hayes 1981). Moreover, SSp activation can produce both excitatory and inhibitory effects in the cuneate (Jabbur and Towe 1960, Andersen, Eccles et al. 1964, Kuypers and Tuerk 1964, Rustioni and Hayes 1981, Cole and Gordon 1992, Canedo, Marino et al. 2000), suggesting that sensory cortex provides top-down regulation of its afferent inputs. We first set out to determine whether this modulation is exclusive to SSp, or if it might involve corticofugal neurons in other cortical regions. Using combinatorial genetic and viral tools, we broadly labeled cortical neurons that project to the cuneate, to cervical spinal cord, or to both. We found that corticospinal neurons throughout contralateral SSp send collateral projections to Cu, whereas corticospinal neurons in primary motor cortex (MOp) and secondary motor cortex (MOs) mostly avoid the core region of Cu (**Suppl. Fig. 8A**). Supporting these findings, targeted anterograde labeling revealed that while SSp densely innervates the core region of Cu, sparse MOp projections are mostly found in ventral cuneate regions (**Suppl. Fig. 8B**), in line with stimulation experiments in rats that found extensive SSp but little MOp excitatory effects in the middle cuneate (Shin and Chapin 1989). We also found corticofugal neurons that project to Cu in a broad, contralateral anterior cortical region that we refer to as rostral sensorimotor cortex (rSM), which do not appear to project to the spinal cord (**Suppl. Fig. 8A**). Together, these results identify candidate circuits throughout cortex for top-down cuneate modulation.

To explore the synaptic connectivity of these corticofugal projection neurons more directly, we used monosynaptic retrograde rabies tracing approaches to identify synaptic inputs to Cu circuits (**Fig. 5A**). First, we found direct cuneolemniscal innervation by corticofugal neurons in SSp as well as from the cervical dorsal root ganglia, as expected. We also found cortical inputs arising from contralateral supplemental somatosensory cortex (SSs), but sparse or absent labelling in MOp or the more anterior rSM region (**Fig. 5A**, left). Next, we reasoned that in addition to directly exciting cuneolemniscal neurons, some descending projections might also inhibit tactile transmission by recruiting the Cu inhibitory circuits that we identified. To test this idea, we restricted rabies tracing to Cu inhibitory neurons in *VGAT-Cre* mice (**Fig. 5A**, right). As with cuneolemniscal tracing, we found that cortical neurons in SSp and SSs also innervate these inhibitory circuits, providing a potential explanation for previous findings that sensory cortex can both excite and inhibit cuneate neurons. Cervical dorsal root ganglia neurons also target these inhibitory neurons, suggesting that sensory pathways can drive feed-forward inhibition of tactile circuits, potentially as a means for decorrelating responses to segregate inputs or sharpen receptive fields (Soto, Aguilar et al. 2004, Witham and Baker 2011, Jorntell, Bengtsson et al. 2014). Most strikingly, we also found a much larger population of previously unidentified rSM neurons that directly innervate Cu inhibitory neurons, but do not target cuneolemniscal neurons (**Fig. 5A**, right).

**Figure 5.**
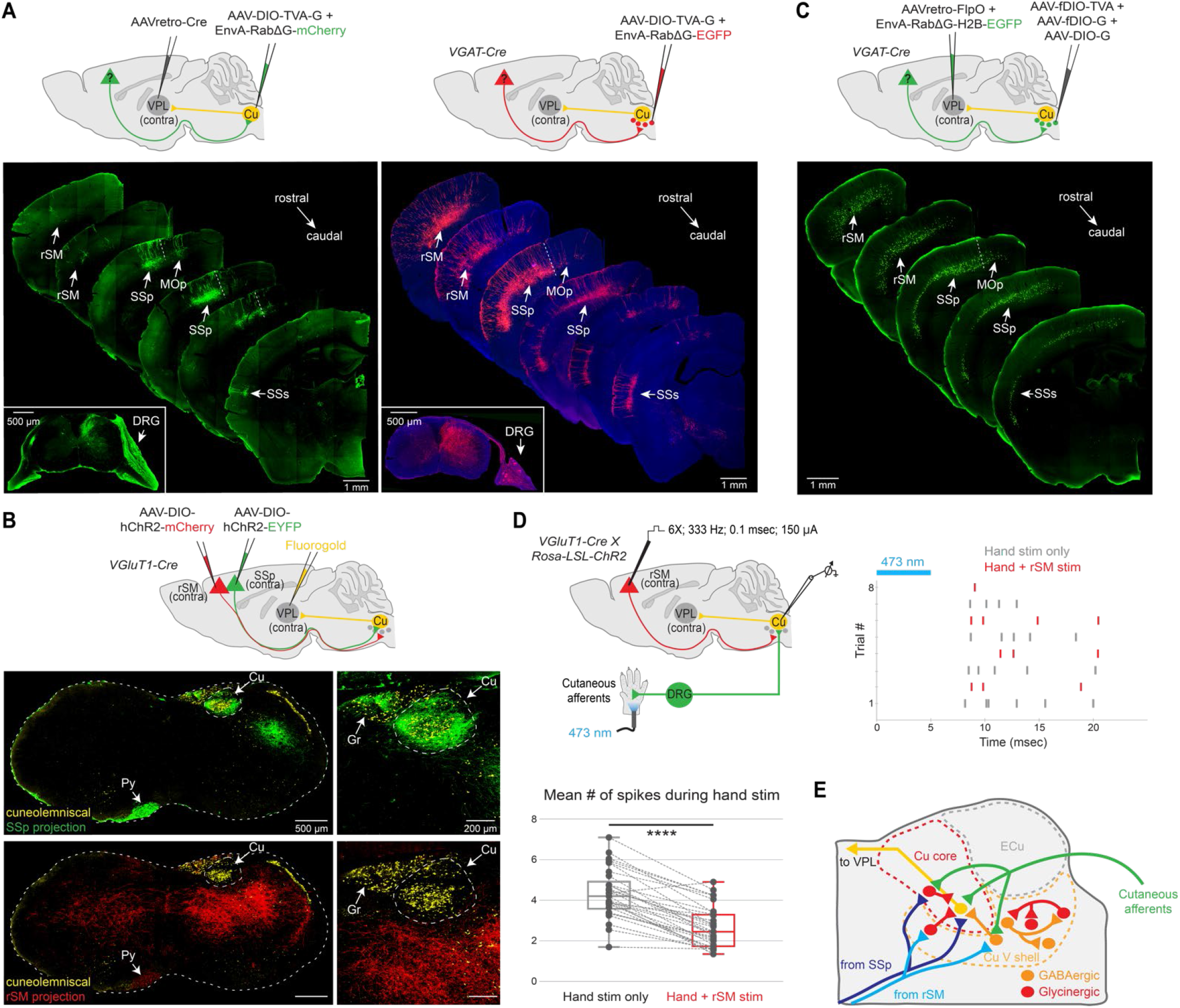
Distinct corticofugal pathways target cuneate circuits. **(A)** Left: Monosynaptic retrograde rabies tracing from cuneolemniscal (CL) neurons (3 mice). Cortical inputs arise almost exclusively from contralateral primary (SSp) and supplemental (SSs) somatosensory cortices, with sparse or absent labelling in primary motor cortex (MOp) or rostral sensorimotor cortex (rSM). Ipsilateral DRG neurons (C7) were also labeled (inset). Right: Monosynaptic retrograde rabies tracing from Cu inhibitory neurons in *VGAT-Cre* mice (3 mice). Cortical inputs arise from contralateral SSp and SSs, but also include a large population of corticofugal neurons throughout rSM. Ipsilateral DRG neurons (C7) were also labeled (inset). (See **Suppl.** Fig. 9 for equivalent results with complementary viral approaches). **(B)** Dual viral anterograde tracing of corticofugal projections in the same *VGluT1-Cre* mice from contralateral SSp (green) and rSM (red), with retrograde Fluorogold labeling of CL neurons (yellow; 2 mice). SSp and rSM corticofugal projections are largely non-overlapping. SSp axons densely innervate the core region of the contralateral Cu and Gr, where cuneolemniscal neurons are located, while rSM axons (red) are excluded from Cu and Gr core regions and innervate more ventral brainstem, including the Cu ventral shell region where inhibitory neurons targeting the Cu core are located. Both corticofugal populations descend through the ipsilateral pyramidal tract (Py) and reach the contralateral brainstem via the pyramidal decussation. Viruses were switched across mice with no change in results (see **Suppl. Fig. 8** for equivalent results with complementary viral approaches). **(C)** Left: Disynaptic retrograde rabies tracing (nuclear localized) from CL neurons, through presynaptic Cu inhibitory neurons, to corticofugal inputs (4 mice). Mirroring the results of monosynaptic tracing from Cu inhibitory neurons ((A), right), cortical inputs arise from contralateral SSp, SSs, and throughout rSM, revealing a feed-forward inhibitory link from these corticofugal populations to CL neurons (see **Suppl. Fig. 9C** for control experiments). **(D)** *In vivo* extracellular recording of tactile-responsive units in Cu while stimulating rSM. Near threshold tactile responses in Cu were elicited by photostimulating ChR2-expressing afferents in the pad of the ipsilateral hand of *VGluT1-Cre* mice (top left, 5 msec). Example spike raster plots (top right) from one recording site across interleaved trials show that tactile-evoked spikes in Cu are suppressed when rSM is electrically stimulated (red; 6 pulses, 333 Hz, 0.1 msec duration, 150 µA; first pulse 30 msec before tactile stimulation at time 0). Quantification across all recordings (bottom left; 33 recordings across 3 mice; 25 msec time window from hand stimulation; \*\*\*\**P* < 0.0001; paired t test). **(E)** Schematic of local and long-Cu connections (see **Suppl. Fig. 10**).

Supporting these findings, dual anterograde labeling of corticofugal neurons in SSp and rSM in *VGluT1-Cre* mice showed essentially non-overlapping and complementary projection patterns; SSp heavily innervates the Cu core region, where cuneolemniscal neurons reside, while rSM innervates more ventral brainstem, including the Cu ventral shell region where inhibitory neurons are located, but completely avoids the Cu core (**Fig. 5B**). Additional combinatorial retrograde viral targeting from the cervical spinal cord confirmed that SSp projections to the Cu core are indeed collaterals of corticospinal projections (**Suppl. Fig. 8C**). In contrast, rSM projections also descend through the pyramidal tract and decussate, but instead innervate the Cu ventral shell region and do not send projections to forelimb regions of the spinal cord (**Suppl. Fig. 8D**). Finally, for a more comprehensive identification of regions that innervate Cu circuits, we used complementary rabies tracing approaches with a nuclear-localized fluorophore and automated serial two-photon tomography to quantify neuronal populations throughout the brain that innervate either cuneolemniscal or Cu inhibitory neurons. This labeling confirmed our retrograde and anterograde anatomical analysis and also identified several subcortical regions as candidate modulators of signaling in the dorsal column nuclei (**Suppl. Tables 1,2** and **Suppl. Fig. 9A,B**). Together, these experiments define the descending pathways capable of modulating tactile feedback through direct excitation or feed-forward inhibition of cuneolemniscal neurons.

Corticofugal neurons in rSM project to a relatively large region of the brainstem (**Fig. 5B** and **Suppl. Fig. 8D**), leaving open the possibility that the inhibitory neurons they innervate (**Fig. 5A**, right) are not the same neurons that modulate tactile transmission within Cu. To explore this question, we designed a disynaptic rabies tracing approach to determine whether rSM corticofugal neurons target inhibitory neurons that then target cuneolemniscal neurons. First, as a control, we initiated monosynaptic rabies tracing selectively from cuneolemniscal neurons that target VPL thalamus, finding the expected inputs from SSp, cervical dorsal root ganglia, and local Cu inhibitory neurons (**Suppl. Fig. 9C**). In a separate group of mice, we again initiated monosynaptic tracing from cuneolemniscal neurons while introducing supplemental rabies G-protein expression selectively in Cu inhibitory neurons in *VGAT-Cre* mice, thereby enabling disynaptic rabies virus spread only through the inhibitory circuits that innervate the cuneolemniscal starter population (**Fig. 5C**). Consistent with our previous anatomical results (**Fig. 5A,B** and **Suppl. Fig. 9A,B**), we found broad labeling in rSM as well as SSp, demonstrating a feed-forward inhibitory link from these corticofugal populations to cuneolemniscal neurons (**Fig. 5C**).

Finally, we wanted to determine whether this rSM corticocuneate projection can indeed inhibit tactile feedback within Cu. Motivated by work suggesting that descending modulation of tactile feedback is most apparent when inputs are not saturating (Towe and Jabbur 1961), we developed a recording approach where we could more flexibly adjust stimulus strength during each recording. Using a genetic approach to express the excitatory opsin ChR2 in sensory afferents of *VGluT1-Cre* mice, we recorded *in vivo* extracellular responses in the Cu of anesthetized mice while reducing the intensity of photostimulation on the ipsilateral pad of the hand to near spiking threshold (**Fig. 5D**). We then delivered electrical stimulation to contralateral rSM just before tactile photoactivation and found a consistent suppression of tactile-evoked spikes in Cu (38.56% ± 2.97% SEM reduction in spike number; **Fig. 5D**). Together, these results demonstrate that rSM corticofugal neurons can elicit inhibitory modulation of tactile feedback, revealing a newly defined, top-down cortical pathway for regulating ascending somatosensory information (**Fig. 5E** and **Suppl. Fig. 10**).

## Discussion

Focusing on tactile signaling in the cuneate nucleus, we have established the functional connectivity of local GABAergic and glycinergic circuits that target cuneolemniscal neurons and enable bidirectional modulation of somatosensory feedback as it enters the brain. Perturbing the activity of these inhibitory circuits can suppress or enhance tactile responses in the cuneate and disrupts the performance of behaviors that rely on sensory feedback from the hand. Distinct top-down neocortical circuits show complementary projection patterns that together provide circuit mechanisms for excitation or feed-forward inhibition of cuneolemniscal transmission to the thalamus (**Fig. 5E** and **Suppl. Fig. 10**). These results uncover new anatomical and functional circuit architecture for the adjustment of tactile feedback critical for the execution of dexterous forelimb behaviors. More broadly, these findings provide insight into general circuit mechanisms, analogous to those identified for other sensory pathways (Sillito, Jones et al. 1994, Gilbert and Sigman 2007, Lee, Carvell et al. 2008, Fink, Croce et al. 2014, McComas 2016, Confais, Kim et al. 2017, Liu, Latremoliere et al. 2018, Schneider, Sundararajan et al. 2018), that can attenuate disruptive feedback to facilitate successful behavior.

There are several reasons why suppression of tactile signals in the cuneate would be beneficial. Center-surround inhibition could sharpen receptive fields to augment resolution and acuity (Canedo 1997, Soto, Aguilar et al. 2004, Witham and Baker 2011). In addition, feedback caused by movement should be attenuated to counteract disruptive feedback delays and to distinguish expected self-generated reafference from unexpected exafferent signals (Coulter 1974, Chapman 1994, Blakemore, Frith et al. 1999, McComas 2016, Azim and Seki 2019). Conversely, amplification of tactile feedback could, in principle, improve discrimination and select the inputs most relevant to a behavior (Chapman 1994, Canedo 1997, Canedo, Marino et al. 2000, Azim and Seki 2019). Moreover, excitation could convey predictions of upcoming events during movement, preparing sensory circuits to process impending feedback through anticipatory modulation (Kuypers and Tuerk 1964, Wall 1970, Johansson and Flanagan 2009). Given the utility of bidirectional modulation, attenuation and augmentation are likely to occur simultaneously, as is seen with the coincident suppression of cutaneous and enhancement of proprioceptive feedback in the spinal cord during wrist movement (Confais, Kim et al. 2017).

The circuits described here could provide the means to regulate distinct channels of tactile information as they ascend into the brain. The large majority of corticofugal neurons we identified target inhibitory cuneate neurons, suggesting that widespread and perhaps somewhat indiscriminate inhibition is desirable, or at least necessary, for coordinated behavior (Chapin and Woodward 1981, McComas 2016). Indeed, broad suppression of somatosensory feedback and an increase in psychophysical detection thresholds before and during movement are well documented (Ghez and Pisa 1972, Coulter 1974, Chapin and Woodward 1981, Milne, Aniss et al. 1988, Blakemore, Frith et al. 1999). One might speculate that preparatory activity in corticofugal neurons, including those across the large region of rostral cortex that we describe, could mediate this movement-related suppression. In contrast, direct innervation of cuneolemniscal neurons was found almost exclusively from excitatory neurons in primary somatosensory cortex, which is the cortical region that serves as the main recipient of cuneolemniscal signals. There is evidence that cortical connectivity to the cuneate is somatotopically aligned (Cheema, Whitsel et al. 1983, Aguilar, Rivadulla et al. 2003, Loutit, Vickery et al. 2020), suggesting that in the face of broad suppression, specific cortical recipients might select and augment the feedback they receive to facilitate ongoing and future movements (Wall 1970, Canedo 1997, Canedo, Marino et al. 2000). A similar feed-forward facilitation might be at play in the spinal cord, where somatosensory corticospinal neurons that are responsive to light touch sensitize subsequent cutaneous transmission from spinal interneurons (Liu, Latremoliere et al. 2018). The behavioral implications of cortical inhibition and excitation of cuneate signaling remain difficult to define, in part because the corticofugal pathways we describe collateralize to many targets as they descend through the pyramidal tract (Rustioni and Hayes 1981). Reliable approaches to perturb specific axon collaterals while leaving other targets unaffected will be needed to address whether broad attenuation punctuated by selective activation characterizes tactile processing in the cuneate during movement. Moreover, top-down modulation of cuneate might have widespread effects. Cuneate neurons target diverse structures, including the cerebellum, pontine nucleus, inferior olive, red nucleus, and superior colliculus (Berkley, Budell et al. 1986, Loutit, Vickery et al. 2020). Whether these distinct cuneate output channels are differentially modulated remains unknown, but this scenario could provide added flexibility, allowing cortical and subcortical areas to receive different versions of the same peripheral signals.

The circuit organization we find is consistent with a model in which top-down pathways convey sensory predictions of bottom-up sensory signals (Adams, Shipp et al. 2013). One formulation of this predictive processing framework is that prediction error neurons sit at the interface of ascending and descending pathways and come in two flavors: positive prediction error neurons that signal unexpected inputs, and negative prediction error neurons that respond to the absence of a predicted input (Keller and Mrsic-Flogel 2018). A putative circuit implementation takes the form of a positive prediction error neuron receiving bottom-up excitation and top-down feed-forward inhibition, while a negative prediction error neuron receives the reverse – when excitation exceeds inhibition in either class of neuron, a corresponding error signal is sent to update an internal model and modify future predictions (Keller and Mrsic-Flogel 2018). In principle, these types of error predictions could emerge at any, or every, layer of the sensorimotor hierarchy. Indeed, at the first layer of tactile processing in the brain, we find bottom-up sensory and top-down cortical pathways forming all of the requisite excitatory and feed-forward inhibitory connections (**Fig. 5E** and **Suppl. Fig. 10**). A challenge will be to establish the resolution needed to determine whether each of the circuit elements match up in the appropriate combinations. Alongside advancing technologies for cuneate recording in behaving animals (Suresh, Winberry et al. 2017), these approaches should help to resolve whether the connectivity and recruitment of neurons in the dorsal column pathway are reflective of hierarchical predictive processing.

## Acknowledgements

We are grateful to Phong Nguyen and Graham Salmun (Salk Institute) for assistance with mouse husbandry, histology, and lab operations; Byungkook Lim (University of California, San Diego) for the AAV-DIO-mRuby-T2A-Syp-EGFP plasmid; Troy Margrie (Sainsbury Wellcome Centre) for the AAV-fDIO-N2cG-H2B-GFP plasmid; Ali Cetin (Allen Institute for Brain Science) for the EnvA-RabΔG-H2B-EGFP plasmid; Adam Hantman (Janelia Research Campus) for the *VGluT1-Cre*, *Gad2-Flp*, and *GlyT2-Flp* mouse lines; Frank Cardone, Mark Stambaugh, Jeff Sandubrae (Qualcomm Institute, University of California, San Diego), and Dan Butler (Salk Institute) for assistance developing the tactile orienting assay; and Denise Ramirez (Whole Brain Microscopy Facility, University of Texas Southwestern Medical Center) for assistance with serial two-photon tomography and quantification. We thank Sho Aoki, Andrew Fink, Kee Wui Huang, Denis Jabaudon, Kazuhiko Seki, John Tuthill, Michael Yartsev, and members of the Azim lab for valuable discussion and comments on the manuscript.

## Author Contributions

J.M.C. and E.A. initiated the project and J.M.C., A.B., M.I., and E.A. designed the experiments. J.M.C. performed viral injections, anatomical experiments, and *in vivo* electrophysiological recordings and J.M.C., J.T., and E.A. analyzed the data. A.B. performed slice electrophysiology experiments and A.B., M.I., and E.A. analyzed the data. N.B. performed string pull behavioral experiments and N.B., J.T., and E.A. analyzed the data. A.B., M.I., and E.A. designed the tactile orienting assay, M.I. performed the experiments, and M.I., J.T., and E.A. analyzed the data. E.A., J.M.C., A.B., M.I., J.T. and N.B. prepared figures and Methods text. E.A. wrote the manuscript with input from all authors.

## Data and materials availability

All data are in the main text, supplementary materials, or are available from the corresponding author upon request. Analysis code is available from the corresponding author upon request. Design and code for the tactile orienting assay are available at www.github.com/azimlabsalk.

## Materials and Methods

### Mice

Procedures performed in this study were conducted according to US National Institutes of Health guidelines for animal research and were approved by the Institutional Animal Care and Use Committee of The Salk Institute for Biological Studies. Approximately equal numbers of adult male and female mice were used for all experiments and data were combined because no sex differences were observed. Injections for tracing ascending sensory afferents from peripheral targets were performed on postnatal day (P)6-P9 pups, and tissue was collected after P56. All mice were maintained on a C57BL/6 background and housed on a 12:12 hour light cycle.

The following mouse lines were used: Wild-type (**Figs. 1A-F, 2A-C, 5A**, **Suppl. Figs. 2, 6, 7D**, **8B-D**, and **9C**; The Jackson Laboratory and in-house colony); *Avil-Cre* (**Suppl. Fig. 1A**; B6.129P2-*Avil^tm2(cre)Fawa^*/J; The Jackson Laboratory, 032536); *PV-Cre* (**Suppl. Fig. 1B**; B6.129P2-*Pvalb^tm1(cre)Arbr^*/J; The Jackson Laboratory, 017320); *VGAT-Cre* (**Figs. 1G-I, 2D-G, 3, 4, 5A,C**, **Suppl. Figs. 4**, **5, 7B,C**, **9B, Supplementary Movies 1-3,** and **Suppl. Table 2**; B6J.129S6(FVB)-*^Slc32a1tm2(cre)Lowl^*/MwarJ; The Jackson Laboratory, 028862); *GAD1-EGFP* (Tamamaki, Yanagawa et al. 2003) (**Suppl. Fig. 3A**); *GlyT2-EGFP* (Zeilhofer, Studler et al. 2005) (**Suppl. Fig. 3B**); *GAD2-FlpO* (Alhadeff, Su et al. 2018) (**Suppl. Fig. 3C**; Hantman, Janelia Research Campus); *GlyT2-FlpO* (**Suppl. Fig. 3D**; Hantman, Janelia Research Campus); *VGluT1-Cre* (Huang, Sugino et al. 2013) (**Fig. 5B,D;** *Slc17a7-IRES-Cre*; Hantman, Janelia Research Campus); *VGluT2-Cre* (**Suppl. Fig. 9A** and **Suppl. Table 1**; B6J.129S6(FVB)-*Slc17a6^tm2(cre)Lowl^*/MwarJ; The Jackson Laboratory, 028863); *Rosa-LSL-tdTom* (**Suppl. Fig. 8A**; Ai14; B6.Cg-*Gt(ROSA)26Sor^tm14(CAG-tdTomato)Hze^*/J, The Jackson Laboratory, 007908); *Rosa-FSF-tdTom* (**Suppl. Fig. 3C,D**; Ai65F; B6.Cg-*Gt(ROSA)26Sor^tm65.2(CAG-tdTomato)Hze^*/J; The Jackson Laboratory, 032864); *Rosa-LSL-ChR2* (**Fig. 5D**; Ai32; B6;129S-*Gt(ROSA)26Sor^tm32(CAG-COP4*H134R/EYFP)Hze^*/J, The Jackson Laboratory, 012569).

### Viruses

The following adeno associated viruses (AAVs) were used, with serotype and titer (vg/ml) indicated: AAV5-EF1a-DIO-hChR2(H134R)-EYFP (**Suppl. Fig. 1A**; Penn Vector Core; 7.4 x 10^12^; Addgene plasmid #20298); AAV5-CMV-EGFP (**Suppl. Figs. 1A, 6** and **7D**; Salk Vector Core; 2.08 x 10^12^; Addgene plasmid #32395); AAV9-FLEX-rev-ChR2-tdTomato (**Suppl. Figs. 1B** and **8B**; Penn Vector Core; 1.4 x 10^13^ for muscle injections; else used at 2.8 x 10^12;^ Addgene plasmid #18917); AAV9-CAG-FLEX-EGFP (**Suppl. Fig. 1B;** Salk Vector Core; 1.4 x 10^12^; Addgene plasmid #51502); AAV1-hSyn-DIO-mRuby2-T2A-Synaptophysin-EGFP (Knowland, Lilascharoen et al. 2017) (**Fig. 1G**; Vigene Biosciences; 2.5 x 10^12^; Lim, UCSD); AAV5-hSyn-DIO-oChIEF-Citrine (**Figs. 1H,I, 2D,E, 3, 4, Suppl. Figs. 4A-C, 5,** and **7B,C, and Supplementary Movies 1,2**; Salk Vector Core; 2.5 x 10^12^; Addgene plasmid #50973); AAV1-hSyn-SIO-stGtACR2-FusionRed (**Fig. 2F,G** and **Supplementary Movie 3**; Vigene Biosciences; 2.5 x 10^12^; Addgene plasmid #105677); AAV1-hSyn-DIO-TVA66T-tdTomato-CVS-N2cG (**Fig. 5A**, **Suppl. Figs. 3C,D, 9A,B,** and **Suppl. Tables 1,2;** Columbia Vector Core; 2.5 x 10^12^); AAV2-retro-EF1a-Cre (Tervo, Hwang et al. 2016) (**Fig. 5A**, **Suppl. Figs. 3C,D** and **8;** Salk Vector Core; 1.0 x 10^12^; Addgene plasmid #55636); AAV1-SynP-DIO-splitTVA-EGFP-B19G (**Fig. 5A**; UNC Vector Core; 3.9 x 10^12^; Addgene plasmid #52473); AAV2-retro-CMV-EGFP (**Suppl. Fig. 8A**; Salk Vector Core; 3.7 x 10^12^; Addgene plasmid #32395); AAV1-EF1a-DIO-hChR2(H134R)-EYFP (**Fig. 5B** and **Suppl. Fig. 8B-D**; UNC Vector Core; 4.0 x 10^12^ and Penn Vector Core; 4.25 x 10^12^; Addgene plasmid #20298); AAV1-EF1a-DIO-hChR2(H134R)-mCherry (**Fig. 5B** and **Suppl. Fig. 8B**; Salk Vector Core; 4.0 x 10^12^; Addgene plasmid #20297); AAV2-retro-Ef1a-FlpO (**Fig. 5C** and **Suppl. Fig. 9C**; Salk Vector Core; 2.5 x 10^12^; Addgene plasmid #55637); AAV8-CAG-fDIO-TC (TVA-mCherry) (**Fig. 5C** and **Suppl. Fig. 9C**; Salk Vector Core; 1.25 x 10^12^; Addgene plasmid #67827); AAV1-Syn-fDIO-N2cG-H2B-GFP (**Fig. 5C** and **Suppl. Fig. 9C**; Vigene Biosciences; 2.5 x 10^12^; Margrie, Sainsbury Wellcome Centre); AAV1-CAGGS-DIO-H2B-GFP-P2A-N2cG (**Fig. 5C;** Salk Vector Core; 2.5 x 10^12^; Addgene plasmid #73475). The following rabies viruses were used: EnvA-Rab-CVS-N2cΔG-EGFP (**Fig. 5A** and **Suppl. Fig. 3C,D**; Columbia Vector Core; 1.0 x 10^9^; Addgene plasmid #73461); EnvA-Rab-pSADΔG-mCherry (**Fig. 5A**, Salk Vector Core; 1.0 x 10^9^; Addgene plasmid #32636); EnvA-Rab-CVS-N2cΔG-H2B-EGFP (**Fig. 5C**, **Suppl. Fig. 9**, and **Suppl. Tables 1,2;** Salk Vector Core; 6.4 x 10^9^; Cetin, Allen Institute for Brain Science).

### Antibodies and tracers

The following primary antibodies were used: goat anti-cholera toxin B subunit (1:2000; List Biological Laboratories, #703); rabbit anti-GFP (used for EGFP, citrine, EYFP; 1:1000; Thermo Fisher Scientific, A-11122); goat anti-GFP (used for EGFP, citrine, EYFP; 1:1000; Abcam, ab6673); rabbit Living Colors anti-DsRed (used for mCherry, tdTomato; 1:1000; Takara Bio, 632496); goat anti-RFP (used for mCherry, tdTomato; 1:1000; Sicgen, AB1140-100); rabbit anti-Fluorogold (1:500; Millipore Sigma, AB153-I). The following conjugated secondary antibodies were used at a concentration of 1:1000: donkey anti-goat-488 (Jackson ImmunoResearch Laboratories, 705-545-147); donkey anti-goat-555 (Thermo Fisher Scientific, A-21432); donkey anti-rabbit-488 (Jackson ImmunoResearch Laboratories, 711-545-152); donkey anti-rabbit-555 (Thermo Fisher Scientific, A-31572); donkey anti-rabbit-647 (Jackson ImmunoResearch Laboratories, 711-605-152); biotin-SP donkey anti-goat (Jackson ImmunoResearch Laboratories, 705-065-147); biotin-SP donkey anti-rabbit (Jackson ImmunoResearch Laboratories, 711-065-152). The following fluorophore conjugated streptavidin complexes were used: streptavidin-488 (Thermo Fisher Scientific, S11223); streptavidin-555 (Thermo Fisher Scientific, S32355); streptavidin-647 (Thermo Fisher Scientific, S21374). The following retrograde tracers were used: Fluorogold (4% solution in water; Fluorochrome); red retrobeads (Lumafluor); unconjugated cholera toxin B subunit (CTB; 1% in 0.1M phosphate buffer; List Biological Laboratories, #104).

### Immunohistochemistry and imaging

Animals used for histological purposes were perfused with 10 ml cold PBS and 20-25 ml cold 4% paraformaldehyde (PFA) in 0.1M phosphate buffer. Brains, spinal cords, and DRGs were removed and postfixed overnight at 4°C in 4% PFA in 0.1M phosphate buffer. Tissues were then transferred to a 30% sucrose solution for a minimum of 48 hours before being sectioned on a sliding microtome at 40 µm thickness. In most cases, native fluorescence was amplified by immunohistochemistry. For 2-way signal amplification, sections were washed in PBS, incubated for 20 min in PBS + 0.25% Triton X-100, and blocked in PBS + 5% donkey serum for 60 min. Sections were incubated for 1-3 days at 4°C in primary antibodies diluted in PBS + 0.25% Triton X-100 + 5% donkey serum at the concentrations listed above. Following primary antibody incubation, sections were washed 3-5 times in PBS and incubated overnight at 4°C with conjugated secondary antibodies at the concentrations listed above. Sections were then washed 5 times in PBS, mounted onto glass slides and coverslipped with Mowiol mounting media (Cold Spring Harbor Protocols). In cases where axonal fibers were being visualized or additional signal amplification was required, a modified tyramide signal amplification (TSA) protocol was used (Adams 1992). For TSA amplification, sections were washed in PBS, incubated for 20 min in PBS + 0.25% Triton X-100, quenched for 30 min in PBS containing 0.6% H_2_O_2_, and blocked for 60 min in PBS + 5% donkey serum. Sections were then incubated for 2-3 days in primary antibody solution, which was diluted an additional 5-fold from the concentrations listed above. Sections were washed thoroughly in PBS and then incubated overnight with a solution containing biotinylated secondary antibodies at concentrations listed above. Sections were then washed and incubated sequentially with an avidin-biotin complex for 30 min (ABC kit; Vector Laboratories) and with biotinyl tyramide (Adams 1992) (1:2500 dilution for 30 min). Sections were washed thoroughly and incubated overnight with conjugated streptavidin at the concentrations listed above. Sections were again washed thoroughly, mounted onto glass slides and coverslipped with Mowiol mounting media. Processed sections were imaged on either a slide scanner (Olympus VS120) at 10X magnification or on a confocal microscope (Zeiss LSM 700; various magnifications). A motorized stage was used to facilitate the creation of montage images and z-axis image stacks. Images were post-processed in Photoshop and Illustrator (Adobe).

### Surgical procedures

#### General

Surgical procedures performed on adult mice were carried out under isoflurane anesthesia (1-3%). The surgical site was shaved and cleaned with betadine and alcohol and animals were given subcutaneous injections of carprofen (5 mg/kg) and bupivacaine (2 mg/kg). Throughout all surgical procedures, animals were kept on a heating pad to maintain body temperature and eye lubricant was applied. Viral injections were performed using pulled glass capillaries and a Nanoject III (Drummond Scientific) mounted to a stereotaxic manipulator. For cuneate targeting, animals were placed in a stereotaxic frame equipped with custom-made ear bars to allow the head to be dorsiflexed 30° from horizontal. From this position, an incision was made at the back of the head and the skin and overlying muscles were retracted to expose obex. In some cases, the most caudal portion of the occipital bone was removed to permit access to more rostral aspects of the cuneate nucleus. Coordinates were determined relative to obex and the dura was pierced with a 30½ gauge syringe to facilitate inserting the glass capillary. For cortical and thalamic injections, animals were placed in a stereotaxic frame in a flat skull position. A skin incision was made at the top of the head, coordinates were determined relative to bregma, and a small craniotomy was made with a dental drill in the skull overlying the injection target. Headplate fixation was performed in a similar fashion, leaving the skull intact. For spinal cord and DRG injections, animals were placed in a flat skull position and the spine was stabilized by securing the tail of the animal with a retractor positioned to produce slight tension along the antero-posterior axis. An incision was made over the cervical spinal cord, the skin and overlying muscles were retracted exposing the vertebrae, muscle was cleaned from the spinal cord using the back of a scalpel blade and fine cotton swabs, and a laminectomy of the cervical (C)6 or C7 vertebrae was performed to improve access to the spinal cord. Injections were performed by piercing the dura with a 30½ syringe to facilitate penetration of the glass capillary into the spinal cord tissue. For all surgical procedures, dorso-ventral coordinates were measured relative to the surface of the brain or spinal cord. All viruses were typically injected at a rate of 3-5 nl/sec. Following all surgical procedures, overlying muscle layers and skin were sutured (6-0 black braided silk, Ethicon), animals were given a subcutaneous injection of buprenex-SR (1 mg/kg), and were allowed to recover on a heating pad. All postoperative animals were housed individually until the end of the experiment.

#### Labeling sensory afferents

a) <u>DRG labeling</u>: To label sensory afferents through viral targeting of the DRG, adult *Avil-Cre* mice were anesthetized and the cervical spinal cord was exposed. After retracting overlying muscles, a laminectomy was performed to remove the C7 vertebra, extending the laminectomy laterally to partially remove the facet joint. A pulled glass capillary was loaded with AAV5-EF1a-DIO-hChR2(H134R)-EYFP and angled outward ∼20°. The capillary was inserted under the remaining facet joint and guided under visual control into the DRG, and a total of 30 nl of virus was injected into the DRG. Overlying muscles and skin were sutured and animals were allowed to survive for 3-4 weeks before being perfused and processed for histological analysis. A second approach to label DRG afferents used injections of AAV5-CMV-EGFP into the DRG of adult wild-type mice (data not shown). Similar results were observed with both approaches.

b) <u>Afferent labeling</u>: To label sensory afferents from their peripheral targets, unconjugated CTB was injected as a tracer. To label cutaneous afferents, P6-P9 mice were anesthetized with isoflurane and 0.5-1.0 µl of 1% CTB solution was injected into the glabrous pad of the hand at a rate of 1 µl/min using a syringe pump (NE-300, New Era Pump Systems) and a Hamilton 10 µl syringe attached to a pulled glass capillary. To label proprioceptive afferents, P6-P9 pups were anesthetized with isoflurane, a small incision was made in the skin overlying the biceps and triceps muscles, and 0.5-1.0 µl of 1% CTB solution was injected into each muscle at a rate of 1 µl/min. Following muscle injections, the capillary was retracted and the skin incision was closed with Vetbond tissue adhesive (3M). Pups were placed on a heating pad until fully recovered before returning to their home cages with their mother. Animals matured to adulthood (∼8 weeks) before being perfused and processed for histology.

c) <u>Proprioceptor labeling</u>: To genetically restrict viral targeting to proprioceptive afferents (Zhang, Morita et al. 1990, de Nooij, Doobar et al. 2013), P6-P9 *PV-Cre* pups were injected with 0.5-1.0 µl of AAV9-FLEX-rev-ChR2-tdTomato or AAV9-CAG-FLEX-EGFP. Virus was injected through a pulled glass capillary into triceps and biceps (1 µl/min) following surgical procedures described above. Following injections, pups were placed on a heating pad until fully recovered before returning to their home cages with their mother. Animals matured to adulthood (∼8 weeks) before being perfused and processed for histology.

#### Targeting local inhibitory circuits within the cuneate region

a) <u>Labeling local inhibitory neurons</u>: To characterize the location of inhibitory neurons targeting the core region of the cuneate, *VGAT-Cre* mice were injected with AAV1-hSyn-DIO-mRuby2-T2A-Synaptophysin-EGFP in the cuneate region at the following coordinates relative to obex: site 1: antero-posterior (A/P) 0.0 mm, medio-lateral (M/L) 0.6 mm, dorso-ventral (D/V) −0.25 mm; site 2: A/P 0.25 mm, M/L 0.8 mm, D/V −0.3 mm; site 3: A/P 0.5 mm, M/L; 1.1 mm, D/V −0.35 mm. A total of 40-50 nl of virus was injected at each site. In the same surgical session, cuneolemniscal neurons were labeled by injecting 4% Fluorogold into the contralateral ventral posterolateral (VPL) thalamus. VPL injections were made at the following locations relative to bregma: site 1: A/P −1.25 mm, M/L 1.75 mm, D/V −3.2 to −3.4 mm; site 2: A/P −1.50 mm, M/L 1.8 mm, D/V −3.3 to −3.5 mm; site 3: A/P −1.85 mm, M/L 1.9 mm, D/V −3.5 to −3.7 mm. A total of 140 nl of Fluorogold solution was injected at each site, with 70 nl injected at the most ventral D/V coordinate and 70 nl injected at the most dorsal coordinate. Animals were left for 3-4 weeks before being perfused and processed for histology.

b) <u>Targeting local inhibitory neurons for electrophysiological recording and perturbation</u>: For slice whole-cell recording, *in vivo* extracellular recording, and behavioral experiments, inhibitory neurons in the cuneate region were targeted by injecting *VGAT-Cre* mice with the following viruses (40-50 nl per site at coordinates described above): AAV5-hSyn-DIO-oChIEF-Citrine and AAV1-hSyn-SIO-stGtACR2-FusionRed. For control behavioral experiments, wild-type mice were injected with AAV5-CMV-EGFP using the same approach. For behavioral experiments, a custom-made fiber optic cannula with a 200 µm core was implanted above the cuneate. Mice were anesthetized and placed in a stereotaxic frame in a flat skull position, and fibers were implanted at the following coordinates relative to lambda: A/P −3.15 mm, M/L 0.8 mm, D/V −4.88 mm. Fibers were cemented to the skull (Tetric EvoFlow, Ivoclar Vivadent) and the skin was sutured around the epoxy. Animals were left for 3-4 weeks before electrophysiological or behavioral experiments. For slice electrophysiology experiments, cuneolemniscal neurons were also labeled by injecting 50-60 nl of red retrobeads into contralateral VPL thalamus (at the sites described above) at least 1 week prior to recording.

c) <u>Targeting GABAergic and glycinergic neurons</u>: *GAD1-EGFP* and *GlyT2-EGFP* mice were used to characterize the regional localization of GABAergic and glycinergic neurons within the cuneate core and ventral shell regions. In these mice, cuneolemniscal neurons were labeled by injecting 1% CTB or 4% Fluorogold into contralateral VPL thalamus (140 nl per site at coordinates described above).

#### Monosynaptic and disynaptic rabies tracing

a) <u>Labeling monosynaptic inputs to cuneolemniscal neurons</u>: To map inputs to cuneolemniscal neurons, a combinatorial viral strategy was used. First, AAV2-retro-EF1a-Cre was injected into contralateral VPL thalamus (50-60 nl per site at coordinates described above) of wild-type mice. Second, Cre-dependent rabies helper virus AAV1-SynP-DIO-splitTVA-EGFP-B19G was injected into the cuneate region (40-50 nl per site at coordinates described above). Animals were left for 3-4 weeks before undergoing an additional surgery where EnvA-Rab-pSADΔG-mCherry was injected into the cuneate region (40-50 nl per site at coordinates described above). Animals were left for an additional 10 days before being perfused and processed for histology.

To determine whether distinct subsets of local inhibitory neurons in the cuneate region (GABAergic and glycinergic) are monosynaptically connected to cuneolemniscal neurons, rabies tracing from cuneolemniscal neurons was carried out as described above in *GAD2-FlpO* x *Rosa-FSF-tdTom* mice or *GlyT2-FlpO* x *Rosa-FSF-tdTom*, enabling rabies labeled cells to be colocalized to fate mapped inhibitory neurons. For this purpose, the low affinity TVA rabies helper virus AAV1-hSyn-DIO-TVA66T-tdTomato-CVS-N2cG was targeted to cuneolemniscal neurons by retrograde viral infection with AAV2-retro-EF1a-Cre from the contralateral VPL thalamus. The TC66T strain of rabies helper contains a mutated low affinity TVA receptor that minimizes nonspecific expression, thus enabling reliable monosynaptic tracing of local connections (Miyamichi, Shlomai-Fuchs et al. 2013). Animals were left for 3-4 weeks before undergoing an additional surgery where EnvA-Rab-CVS-N2cΔG-EGFP was injected into the cuneate. Animals were left for an additional 10 days before being perfused and processed for histology.

To confirm the results of these tracing experiments and to enable automated quantification through nuclear labeling, a second strategy for assessing monosynaptic inputs onto cuneolemniscal neurons was used. First, AAV1-hSyn-DIO-TVA66T-tdTomato-CVS-N2cG was injected into the cuneate region (40-50 nl per site at coordinates described above) of *VGluT2-Cre* mice to express rabies helper constructs only in cuneate excitatory neurons. Animals were then left for 7 weeks before undergoing a second surgery where EnvA-Rab-CVS-N2cΔG-H2B-EGFP (expressing nuclear-localized EGFP) was injected into contralateral VPL thalamus (60-70 nl per site at coordinates described above), thus targeting only the subset of cuneate excitatory neurons that give rise to cuneolemniscal projections. Animals were left for an additional 10 days before being perfused and processed for histology. Similar results were observed with both approaches. To determine the spatial localization and relative size of brain-wide cell populations providing monosynaptic inputs onto cuneolemniscal neurons, labeled brains were processed at the Whole Brain Microscopy Facility at the University of Texas Southwestern Medical Center using a TissueCyte 1000 system (TissueVision). Briefly, this system uses serial tissue sectioning and two-photon tomography to image the entire brain (Ramirez, Ajay et al. 2019). An analysis pipeline was then used to implement supervised machine learning classifiers to generate a fluorescence probability map registered to a standardized 3-D atlas of the adult mouse brain (Common Coordinate Framework v3.0; Allen Institute for Brain Science) (Poinsatte, Betz et al. 2019). Use of the nuclear localized fluorophore aided in segmentation by minimizing errors associated with labeled axons. Ipsilateral refers to the hemisphere containing the rabies virus starter cell population. Data collected includes: region volumes; mean raw intensity of fluorescence in each region across mice; mean intensity of fluorescence per cubic mm in each region; SEM of mean intensity of fluorescence per cubic mm in each region; the mean probability normalized to the highest whole-brain mean intensity per cubic mm; and the SEM of the mean probability normalized to the highest whole-brain mean intensity per cubic mm. Region abbreviations can be found at https://mouse.brain-map.org/static/atlas.

A third strategy for assessing monosynaptic inputs onto cuneolemniscal neurons used an approach similar to the first one but substituted AAV2-retro-Ef1a-FlpO injections into VPL thalamus and Flp-dependent rabies helper viruses (AAV8-CAG-fDIO-TC (TVA-mCherry) and AAV1-Syn-fDIO-N2cG-H2B-GFP) into the cuneate region (see details in control experiment of (c) below). All three monosynaptic tracing approaches labeled indistinguishable cortical populations.

b) <u>Labeling monosynaptic inputs to cuneate inhibitory neurons</u>: To map inputs to cuneate inhibitory neurons, AAV1-hSyn-DIO-TVA66T-tdTomato-CVS-N2cG was injected into the cuneate region of *VGAT-Cre* mice (40-50 nl per site at coordinates described above). Animals were then left for 3-4 weeks before undergoing an additional surgery where EnvA-Rab-CVS-N2cΔG-EGFP was injected into the cuneate region (40-50 nl per site at coordinates described above). Animals were left for an additional 10 days before being perfused and processed for histology.

As above, to confirm the results of these tracing experiments and to enable automated quantification through nuclear labeling, a second strategy for assessing monosynaptic inputs onto cuneate inhibitory neurons was used. First, AAV1-hSyn-DIO-TVA66T-tdTomato-CVS-N2cG was injected into the cuneate region (40-50 nl per site at coordinates described above) of *VGAT-Cre* mice to express rabies helper constructs only in local inhibitory neurons. Animals were then left for 3-4 weeks before undergoing a second surgery where EnvA-Rab-CVS-N2cΔG-H2B-EGFP (expressing nuclear-localized EGFP) was injected into the cuneate region (40-50 nl per site at coordinates described above). Animals were left for an additional 10 days before being perfused and processed for histology. Similar results were observed with both approaches. Quantitative localization of brain-wide cell populations providing monosynaptic inputs onto cuneate inhibitory neurons was performed as described above.

c) <u>Disynaptic tracing of inputs to cuneolemniscal neurons</u>: To determine whether the inhibitory neurons that target cuneolemniscal neurons are the same neuronal population targeted by rSM projections, we used a disynaptic rabies tracing approach. In this paradigm, G-deleted rabies virus tracing was initiated specifically from cuneolemniscal neurons, but local inhibitory cell populations were also selectively supplemented with G-protein (but not with TVA protein). Thus, while the starting cells for rabies virus tracing were restricted to cuneolemniscal neurons, the virus could propagate disynaptically through local inhibitory cells that are synaptically connected to cuneolemniscal starter cells. First, AAV2-retro-Ef1a-FlpO was injected into the contralateral VPL thalamus (60-70 nl per site at coordinates described above) of *VGAT-Cre* mice. Second, Flp-dependent rabies helper viruses AAV8-CAG-fDIO-TC (TVA-mCherry) and AAV1-Syn-fDIO-N2cG-H2B-GFP were injected into the cuneate region (viruses were combined and 40-50 nl of virus solution was injected per site at coordinates described above). In the same surgery, supplemental G protein was delivered specifically to the local inhibitory neurons by injecting the Cre-dependent virus AAV1-CAGGS-DIO-H2B-GFP-P2A-N2cG into the cuneate region (40-50 nl per site at coordinates described above), enabling rabies to jump retrogradely from local inhibitory neurons that are presynaptic to targeted cuneolemniscal cells. Animals were left for 3-4 weeks before undergoing an additional surgery where EnvA-Rab-CVS-N2cΔG-H2B-EGFP (expressing nuclear-localized EGFP) was injected into contralateral VPL thalamus (60-70 nl per site at coordinates described above), selectively targeting cuneolemniscal neurons that express helper viruses. Animals were left for an additional 13 days (to allow for disynaptic spread) before being perfused for histology. As a control, the same procedures were performed in wild-type mice, omitting injection of supplemental G protein (AAV1-CAGGS-DIO-H2B-GFP-P2A-N2cG) into the cuneate region.

#### Labelling cortical axonal projections to the cuneate region

a) <u>Corticofugal projection labeling</u>: Initially we explored which types of corticofugal populations extend axon collateral inputs to the cuneate region. To address this question we used a dual retrograde labeling approach to separately identify corticospinal and corticocuneate populations. First, to label cervical-projecting corticospinal populations, AAV2-retro-EF1a-Cre was injected into the C6-C8 spinal segments at 5 equally spaced A/P locations in *Rosa-LSL-tdTom* reporter mice. At each location, the capillary was lowered to 1.5 mm ventral to the surface of the spinal cord, 25 nl of virus was injected, and the capillary was then raised in discrete 200 µm steps with 25 nl of virus being injected at each step. The most dorsal injection was made 250-300 µm ventral to the surface of the spinal cord. Second, in the same animals AAV2-retro-CMV-EGFP was injected into the cuneate region (40-50 nl per site at coordinates described above) to label all corticocuneate projections, including corticospinal collateral inputs. Animals were left for 3-4 weeks being perfused and processed for histology. A series of 40 µm slices throughout the full extent of the forebrain, spaced 120 µm apart, was immunostained and imaged.

b) <u>SSp and MOp projection labeling</u>: To separately label corticospinal projection neurons in SSp and MOp that send collaterals to the cuneate region, wild-type mice were injected with AAV2-retro-EF1a-Cre into C6-C8 spinal segments, as described above. In half of these mice, AAV9-FLEX-rev-ChR2-tdTomato or AAV1-EF1a-DIO-hChR2(H134R)-mCherry was injected into the contralateral SSp and AAV1-EF1a-DIO-hChR2(H134R)-EYFP was injected into contralateral MOp. In the remaining mice the viruses were switched, with no change in results. The coordinates for each virus injection relative to bregma were as follows: SSp: A/P −0.1 / −0.2 / −0.3 mm, M/L 2.2 / 2.5 mm, D/V −0.6 mm; MOp: A/P 0.2 / 0.4 / 0.6 mm, M/L 1.0 / 1.2 mm, D/V −0.6 mm. At each site, 50 nl virus was injected. Animals were left for 3-4 weeks being perfused and processed for histology.

c) <u>Dual labeling of SSp and rSM projections to the cuneate region</u>: To simultaneously visualize corticofugal projections arising from either SSp or rSM that target the cuneate region, *VGluT1-Cre* mice were injected with AAV1-EF1a-DIO-hChR2(H134R)-EYFP in SSp and AAV1-EF1a-DIO-hChR2(H134R)-mCherry into rSM. In some mice the viruses were switched, with no change in results. For SSp targeting, virus was injected into the cortex at 9-10 equally spaced sites spanning the following coordinates relative to bregma: A/P −1.5 → 0.5 mm, M/L 1.0 → 3.0 mm, D/V −0.4 → −0.7 mm. At each site, 50 nl of virus was injected, spread evenly across the D/V extent. For rSM targeting, virus was injected at the following 7 sites relative to bregma: site 1: A/P 1.7 mm, M/L 1.5 mm, D/V −0.8 mm; site 2: A/P 1.7 mm, M/L 2 mm, D/V −1.0 mm; site 3: A/P 1.7 mm, M/L 2.5 mm, D/V −0.8 to −2.25 mm; site 4: A/P 2.0 mm, M/L 1.5 mm, D/V −0.7 mm; site 5: A/P 2.0 mm, M/L 2.1 mm, D/V −0.7 to −2.25 mm; site 6: A/P 2.25 mm, M/L 1.5 mm, D/V −0.9; site 7: A/P 2.25 mm, M/L 2.0 mm, D/V −0.8 to −1.8 mm. At each site, the pipette was lowered to its most ventral location and 50 nl of virus was injected every 200 µm as the capillary was withdrawn along the D/V axis. In the same surgical session, cuneolemniscal neurons were also labeled by injecting 4% Fluorogold into contralateral VPL thalamus, as described above. Animals were left for 3-4 weeks being perfused and processed for histology.

d) <u>Selective labeling of SSp corticospinal collaterals to the cuneate</u>: To label collaterals arising from corticospinal neurons in SSp, a combinatorial viral strategy was used. First, AAV2-retro-EF1a-Cre was injected into C6-C8 spinal segments, as described above. Second, in the same surgical session, AAV1-EF1a-DIO-hChR2(H134R)-EYFP was injected into contralateral SSp, as described above. Cuneolemniscal neurons were also labeled by injecting 4% Fluorogold into contralateral VPL thalamus, as described above. Animals were left for 3-4 weeks being perfused and processed for histology.

e) <u>Selective labeling of rSM corticofugal collaterals to the cuneate region</u>: To label collaterals arising from corticofugal neurons in rSM, a combinatorial viral strategy was used. First, AAV2-retro-EF1a-Cre was injected into the cuneate region, as described above. Second, in the same surgical session, AAV1-EF1a-DIO-hChR2(H134R)-EYFP was injected into the contralateral rSM, as described above. In some cases, cuneolemniscal neurons were also labeled by injecting 4% Fluorogold into contralateral VPL thalamus, as described above. Animals were left for 3-4 weeks being perfused and processed for histology.

### Slice electrophysiology

#### Slice preparation

Acute brainstem slices containing the cuneate nucleus were prepared from adult (∼8-12 week old) mice. In cases where cuneolemniscal neurons were targeted for whole-cell recordings, animals were injected with red retrobeads into contralateral VPL thalamus at least 1 week prior to recording, as described above. Brain slices were collected, as previously described (Ting, Daigle et al. 2014). Briefly, animals were deeply anaesthetized and decapitated, and the brain was quickly removed and immediately placed in ice-cold cutting solution consisting of (in mM): 92 *N*-methyl-d-glucamine, 30 NaHCO3, 25 D-glucose, 20 HEPES, 1.25 KH2PO4, 5.0 ascorbate, 3.0 pyruvate, 2.0 thiourea, 2.5 KCl, 3.5 MgCl2, and 0.5 CaCl2. Coronal brain slices (250 μm thick) were sectioned using a vibratome (VT1000 S, Leica), and slices were transferred to cutting solution at 37°C for 5-10 min for recovery. Until recording, slices were stored in room temperature holding solution containing (in mM): 92 NaCl, 30 NaHCO3, 25 D-glucose, 20 HEPES, 1.25 KH2PO4, 5.0 ascorbate, 3.0 pyruvate, 2.0 thiourea, 2.5 KCl, 2.0 MgCl2, and 2.0 CaCl2. For recording, slices were transferred to room temperature artificial cerebrospinal fluid (aCSF) containing (in mM): 125 NaCl, 26 NaHCO3, 10 D-glucose, 1.25 NaH2PO4, 3.5 KCl, 2.0 CaCl2, and 2.0 MgCl2. All solutions were continuously oxygenated (95% O2, 5% CO2) at pH 7.4-7.5, with an osmolarity of 290-300 mOsm, adjusted with sucrose as necessary.

#### Whole-cell recording

Individual slices were transferred to a submerged recording chamber mounted on the stage of a fixed-stage microscope (BX51WI, Olympus) equipped with differential interference contrast optics, a 40X water immersion lens, and infrared illumination to view neurons in the slices. The recording chamber was continuously perfused with oxygenated aCSF at room temperature (3 ml/min). Borosilicate patch electrodes were controlled by a motorized micromanipulator (MP-225, Sutter Instrument Company) that had an open tip resistance of 3-6 MΩ when filled with intracellular solution.

For experiments investigating inhibitory activity, the reversal potential of chloride was shifted to ∼0 mV using an intracellular solution containing (in mM): 135 KCl, 10 Hepes, 10 creatine-PO4, 2 Mg-ATP, 0.2 Na-GTP, and 0.5 EGTA, with an adjusted pH of 7.3 and osmolarity of 280-290 mOsm. For all other experiments, the internal solution contained (in mM): 125 K-gluconate, 10 KCl, 10 Hepes, 10 creatine- PO4, 2 Mg-ATP, 0.2 Na-GTP, and 0.5 EGTA, with an adjusted pH of 7.3 and osmolarity of 280-290 mOsm. Tight seals of 1 GΩ or greater were obtained under visual guidance before breaking into whole-cell mode. Neurons expressing fluorescent markers were selected for patching based on their anatomical location. Neuronal health and patch quality were evaluated based on resting membrane potential, input resistance, and responses to depolarizing and hyperpolarizing pulses.

#### Data acquisition and analysis

All results were obtained from cells recorded at least 3 weeks post-virus injection. Whole-cell recordings were obtained using a Multiclamp 700B amplifier (Molecular Devices). Signals were acquired using Clampex software with a Digidata 1550B interface (Molecular Devices). Evoked responses were digitized at 10 kHz, filtered at 2 kHz, and analyzed using Clampfit (version 10.0, Molecular Devices).

Spontaneous events were identified using the event detection feature of Clampfit. Spontaneous event frequency was calculated cumulatively as the total number of events detected over the entire recording period. The instantaneous inter-event interval for spontaneous events was defined as the duration between the onset of consecutive events. To generate the cumulative distribution function of inter-event intervals, all recorded events from all cells were pooled and sorted by increasing duration (x-axis). Interval durations were plotted against the probability (y-axis) that a given inter-event interval would have a nominal value at or below that duration. For evoked events, event latency was defined as the duration from stimulus onset to event onset and averaged over a minimum of 10 trials for each cell. Event jitter was defined as the standard deviation of trial-to-trial event latencies and calculated from a minimum of 10 trials for each cell.

Intrinsic cell properties, including resting membrane potential, input resistance, and action potential threshold, were calculated from traces generated under current-clamp configuration. Resting membrane potential was calculated from at least 5 sweeps as the average membrane potential over a 100 msec period with a 0 pA holding current. Input resistance was calculated with Ohm’s law, using the amplitude of the change in membrane potential in response to a 10-100 pA hyperpolarizing current. Action potential threshold was determined through phase plot of the rate of change of membrane potential versus membrane potential. Depolarizing current was injected incrementally in 10-50 pA steps, and the first evoked action potential was used to calculate action potential kinetics. Synaptic currents were recorded under voltage-clamp configuration to allow for evaluation of kinetics with minimal contamination due to spontaneous fluctuations in membrane potential. The responses of opsin-expressing cells to light pulses were recorded under both current-clamp and voltage-clamp configuration to confirm the presence of light-evoked action potentials and quantify photocurrent amplitudes, respectively.

#### Photostimulation

Photostimulation of opsins was achieved through full-field illumination of the tissue via fluorescent light (X-Cite Turbo; Excelitas Technologies) passed through the microscope objective. Light was passed through a YFP filter (YFP-2427B-000, Semrock; excitation: 500/24-25, emission: 542/27-25) for activation of oChIEF. Unless otherwise noted, a 100 msec light pulse was used for activation with pulse timing triggered by Clampex software.

#### Drugs and drug application

CNQX (10 μM, Abcam) and D-APV (20 μM, Abcam) were used to block AMPA and NMDA receptors, respectively, when quantifying inhibitory inputs. Bicuculline methiodide (10 μM, Abcam) was used to block GABA_A_-mediated synaptic transmission. Strychnine (10 μM, Tocris) was used to block glycine-mediated transmission. All drugs were bath-applied for at least 10 min before assessing post-application responses, with each neuron serving as its own control.

### *In vivo* electrophysiology

#### Animal preparation

Adult mice were anesthetized with isoflurane and placed in a stereotaxic frame (Kopf Instruments) in a flat skull position. The skin and overlying fascia were cut and retracted and the skull was cleaned with cotton swabs and dried. A small burr hole was drilled above the cerebellar vermis and a 125 µm teflon coated tungsten wire (California Fine Wire Company) was lowered ∼1 mm below the surface of the brain and secured in place with epoxy (Tetric EvoFlow, Ivoclar Vivadent), serving as the reference electrode for differential recordings. Next, a headplate was affixed to the skull immediately above bregma with epoxy. The animal was then repositioned in modified ear bars with the head dorsiflexed ∼30° from horizontal and obex and the dorsal brainstem were exposed, as described above. In some cases, the caudal most aspect of the cerebellum was removed by aspiration to improve access to the cuneate. The animal was then removed from isoflurane, administered a cocktail containing 1.2 mg/kg urethane and 20 mg/kg xylazine (Lee and Jones 2018), and transferred to a recording rig where the headplate was clamped with the head dorsiflexed and the cuneate easily accessible. Animals were slightly suspended so that their forelimbs were held in a fixed position above the base of the recording platform. The preamplifier was grounded through an insulated copper strip lined with saline soaked gauze that was wrapped around the tail. The dura and pia overlying the cuneate were removed and the surface of the brainstem was kept moist with a layer of sterile saline or paraffin oil placed in the space created by retracting the muscles. Throughout recordings, animals were kept on a DC-powered heating pad to maintain stable body temperature, and supplemental doses of urethane were given to maintain animals in an areflexic state, assessed by foot pinch.

#### Tactile stimuli

For most experiments, tactile stimuli were applied with 4 small wooden sticks (2.5 mm diameter) that were oriented 90° from one another and rotated across the surface of the pad of the ipsilateral hand using a stepper motor (28BYJ-48m, MikroElektronika) controlled by a microcontroller board (Arduino UNO, Arduino). The board was triggered by a TTL pulse from a Cerebus neural signal processor (Blackrock Microsystems) and was programmed to rotate the stick-wheel at a rate of 60° per sec. There was a 7 sec interval between each complete revolution (1 revolution = 4 tactile stimuli). The duration of each individual stimulus was ∼0.7-0.8 sec and the duration of a full revolution was 6 sec. For each recording site, a total of 20 tactile stimuli were applied (5 revolutions). For optogenetic experiments, 20 light off and 20 light on stimuli were delivered (5 revolutions each), interleaving conditions for each full rotation.

#### Recording

Extracellular recordings were performed using carbon fiber electrodes (∼0.8 MOhm, Carbostar-1, Kation Scientific) mounted to a stereotaxic manipulator (Kopf Instruments) and connected to a Cereplex M headstage (Blackrock Microsystems), which interfaced with the Cerebus neural signal processor. Signals were sampled at 30 kHz and band-pass filtered between 250-5000 Hz. Tactile responsive units in the cuneate were targeted by placing the recording electrode at the following coordinates relative to obex: M/L 0.5 to 0.8 mm, A/P −0.5 to 0.5 mm. The electrode was then slowly lowered as tactile inputs were delivered to the ipsilateral hand and responses were monitored on an oscilloscope and through audio feedback. Robust tactile responses were typically found between 50 and 350 µm below the surface of the brainstem. All data were collected within a 4-6 hour recording session and mice were perfused for histological analysis immediately following the experiment to verify viral expression and electrode placement. All data were analyzed offline by a second experimenter blinded to the experimental conditions. Data consisted of continuous raw data, spiking events (defined as crossing a threshold well above noise), and analog signals marking motor and optogenetic outputs. Cerebus Central Suite (Blackrock Microsystems) or Offline Sorter (Plexon) were used to set event thresholds, and NeuroExplorer (Plexon) was used to export recording files to MATLAB (MathWorks) for further analysis. Due to the high cellular density and anatomical arrangement of tactile responsive clusters within Cu, in some cases the recordings likely reflected multiple responsive units, despite conservative event thresholds. In all cases, comparison of responses was performed from the same recording site, and thus single- or multi-unit recordings from light off and light on conditions were always analyzed in a pairwise fashion.

#### Optogenetic manipulation of cuneate inhibitory neurons

To evaluate the impact of activating local inhibitory circuits on tactile responses in the cuneate, AAV5-hSyn-DIO-oChIEF-Citrine was injected into the cuneate region of adult *VGAT-Cre* mice, as described above. To evaluate the impact of suppressing local inhibitory circuits on tactile responses, AAV1-hSyn-SIO-stGtACR2-FusionRed was injected into the cuneate region of adult *VGAT-Cre* mice, as described above. Following injections, mice were left for at least 3 weeks before electrophysiological recording. For both activation and inactivation, a fiber optic cannula with a 200 µm core (CFM12L10, Thorlabs) was positioned immediately above the cuneate at the surface of the brainstem using a stereotaxic manipulator, and light was delivered using a 470 nm fiber coupled LED (M470F3, Thorlabs) or a 473 nm laser (BL473T8, Shanghai Laser) with an output strength of ∼3-10 mW. At each recording site, optogenetic activation (20 Hz, 5-15 msec pulse width) or inactivation (continuous) occurred on interleaved revolutions of the tactile wheel. Spike thresholds were manually set well above baseline noise during each recording using Cerebus Central Suite. Tactile motor and optogenetic trigger outputs from the Arduino board were sent as analog inputs to the Cerebus neural signal processor for alignment with recording data.

To quantify the effects of optogenetic manipulation, peri-event raster plots were generated and spikes were binned into 100 msec intervals, beginning at the initiation of motor revolution, and continuing for 6 sec, encompassing delivery of 4 tactile stimuli. The number of spikes generated at each recording site across all 20 tactile events (5 revolutions x 4 stimuli) for each condition (light on and light off) were then combined, and bin-averaged histograms with a 100 msec bin size were computed. To separate tactile evoked responses from tactile inter-stimulus interval (ISI) spiking, each 6 sec revolution was segmented into 4 individual 1.5 sec intervals corresponding to a quarter revolution of the stepper motor. Thus, a single tactile stimulus always occurred at some point within each 1.5 sec interval. Within each interval, we estimated the approximate window of tactile stimulation (0.7-0.8 sec) as the period when spike frequency rose to at least 200% above baseline activity, which was usually extremely low. Periods outside of this elevated spike activity were classified as ISI. The process was repeated for each recording site. All responses to optogenetic stimuli were subject to pairwise comparison across light off and light on conditions for each recording site.

#### Assessing rSM corticofugal modulation of tactile responses in the cuneate

To assess the impact of activating descending rSM projections on tactile evoked responses in the cuneate, we developed an approach to deliver near threshold tactile stimuli by photostimulating the pad of the hand of *VGluT1-Cre* x *Rosa-LSL-ChR2* mice with single pulses of 473 nm light (5 msec pulse width, BL473T8, Shanghai Laser). Recordings were made in the cuneate and tactile responsive units were identified, as described above. The laser intensity was adjusted to just above the level required to evoke spiking in the cuneate at each recording site. In addition, bipolar stimulating electrodes, constructed from 125 µm teflon coated tungsten wire (California Fine Wire Company) spaced ∼1 mm apart, were implanted into rSM (see coordinates above). The electrode pair was lowered ∼500-600 µm below the cortical surface and affixed in place with epoxy. In accordance with prior studies (Chambers, Liu et al. 1963, Cole and Gordon 1992), a train of 6 pulses, 0.1 msec duration, at 150 µA was delivered at 333 Hz, with the first pulse delivered 30 msec before optogenetic stimulation of the hand, and 20 trials were performed for each recording site. To quantify the effects of rSM stimulation on tactile-evoked responses, signals were high-pass filtered at 250 Hz, with spike threshold at 133% above baseline noise (calculated using 3-Sigma Peak Heights detector, Offline Sorter, Plexon), and averaged across all recordings per unit or recording site. The total number of light-evoked spikes within 25 msec of the onset of hand optogenetic stimulation was than calculated. All responses were subject to pairwise comparison with and without rSM stimulation for each recording site.

### String pulling behavioral assay

#### Assay

The string pulling task was based on a previously developed assay (Blackwell, Banovetz et al. 2018). Mice were food restricted and maintained at ∼85% of their original body weight. Pre-training for the task was carried out over 2-3 sessions before experimental trials. In the first pre-training session, mice were placed into a box similar to their home cage with 20 cotton strings, 2 mm in diameter, hanging through the top of the cage. Training strings ranged from 30-110 cm in length and were randomly spaced around the box perimeter. Half of the strings were baited with a peanut reward to increase motivation for performing the task. Animals were given 60 min to pull all 20 strings into their cage. If an animal failed to retrieve all 20 strings within the allotted time, the process was repeated the following day. By the end of the second day, all mice had successfully learned to pull the strings into the cage. Animals were then introduced into a 9 x 20 cm plexiglass testing chamber with a pulley system affixed to the top of the front panel. A 70 cm string was routed across the pulley and the end located outside the cage was affixed to a counterweight to encourage animals to pull the entire length of the string without interruption. Animals had to pull the string a total of ∼40 cm to complete the trial, after which a peanut reward was manually dispensed. If the pulling bout was interrupted, the counterweight caused the string to retract to its initial position and mice had to restart the trial. The mice remained in the acquisition box until they successfully performed 4 complete pull trials, at which time they were considered acclimated to task conditions and ready to advance to the video acquisition trials.

For video acquisition trials, mice were placed in the testing chamber and allowed to acclimate before initiating the first trial. For each trial, mice had to retrieve the 70 cm, counterweighted string, after which a peanut reward was manually dispensed. Strings used for video acquisition were marked every 7 mm throughout their length to aid in the analysis of kinematics and pulling performance. Following each successful trial, the mice were given a 45 sec break before initiation of the next trial. Animals typically completed 10-15 trials in a single recording session. Video was acquired at 250 frames/sec by a camera (acA2040, Basler) placed 80 cm from the front panel of the testing chamber using Pylon software (Basler). The camera and testing box remained secured in place, such that the origin of the tracking system remained fixed to an arbitrary point in space.

#### Optogenetic manipulation of cuneate inhibitory neurons

To evaluate the impact of activating local cuneate inhibitory circuits, AAV5-hSyn-DIO-oChIEF-Citrine was injected into the cuneate region of adult *VGAT-Cre* mice, as described above. To evaluate the impact of suppressing local cuneate inhibitory circuits, AAV1-hSyn-SIO-stGtACR2-FusionRed was injected into the cuneate region of adult *VGAT-Cre* mice, as described above. For control experiments, AAV5-CMV-EGFP was injected into the cuneate region of adult wild-type mice, as described above. After viral injections, animals were left for at least 3 weeks before undergoing an additional surgery to implant optical fibers targeting the cuneate, as described above. After fiber implant, animals were returned to their home cage and allowed at least 1 week to recover before behavioral testing.

Mice were briefly placed into an isoflurane induction chamber until they were sedated enough to enable the experimenter to attach a patch cable to the fiber optic cannula. Animals were then placed back in their home cage for 5 min to recover from anesthesia and acclimate to the tethered patch cable, before being transferred to the testing chamber. Mice performed 8-14 trials, interleaving 2 light off trials with 2 light on trials. For all light on trials, a 473 nm laser with output calibrated to ∼13 mW at the end of patch cable was triggered by a programmable Arduino board. For optogenetic activation (oChIEF) experiments, the laser was triggered at the initiation of a pulling bout (10 Hz, 50 msec pulse width) and left on for the duration of the trial. The optimal stimulation pattern was determined empirically by varying stimulation frequency and duration. For optogenetic inactivation (stGtACR2) experiments, the laser was triggered in the middle of the pulling bout (continuous) and left on for the remainder of the trial. Periods of laser activation were marked for video analysis by a low intensity LED placed within the camera field of view but outside the view of the mouse.

#### Data analysis

All string pull data were analyzed by an experimenter blinded to the experimental conditions. Trials were first analyzed by adjusting video playback to 24% real-time, and manually counting the total number of prehension mistakes. Normally, mice alternate pulling the string with the left and right hands in a smooth and rhythmic fashion. Prehension mistakes were defined as any failed attempt to grasp the string on the first attempt, causing a reversal in the direction of hand movement and a second attempt. Multiple grasping attempts in a single cycle were counted as a single prehension mistake. Grasps associated with the initial engagement of the string on the first cycle or at the end of a trial when the string was falling into the cage were excluded from analysis. Occasionally mice grasped the string in their mouths as an alternate strategy for pulling the string downward (most often observed during optogenetic perturbation). In these cases, the first grasp attempt immediately following engagement with the mouth was excluded from analysis. Prehension mistakes were quantified across 5-7 trials per mouse, with a trial defined as a full string pull (∼40 cm).

To quantify string pull kinematics, DeepLabCut (Mathis, Mamidanna et al. 2018, Nath, Mathis et al. 2019) was used to automate tracking of the left and right hands. After tracking, the first ∼7 continuous pulling segments per hand were selected from the longest pulling bout and were defined as a trial, resulting in approximately half of all the pulling bouts being used for kinematic quantification. Files containing x and y pixel coordinates for each hand were imported into MATLAB for kinematic analysis, and quantification was performed across 4-5 trials per mouse. All videos were manually reviewed to identify any outlier coordinates arising from labeling errors, which were infrequent. Any erroneously labeled frames were interpolated and all frame indices were converted to seconds by a conversion factor of 1/250. Next, pixels were converted into distance (mm) using an empirically determined conversion factor; the length of 10 consecutive markings on the string was measured to obtain the average distance between each mark, and pixel distances across each mark were then obtained from 2D image processing software (ImageJ) to compute the conversion factor. These steps were repeated five times and averaged, resulting in a factor of 0.34 mm per pixel. To quantify vertical direction reversals (i.e., the number of times the animal reversed the direction of its hand movement in the y-axis), a peak-to-valley prominence threshold was defined through an empirical process that identified a threshold distance of 4.5 mm as sufficient for detecting nearly all direction reversals. Next, the number of vertical direction reversals exceeding threshold for each hand was calculated as the total number of peaks and troughs. To quantify the vertical pathlength (i.e., the average vertical distance traversed by the hand between direction reversals), the vertical distance between each peak and trough for every direction reversal was quantified and averaged.

### Tactile orienting behavioral assay

#### Assay

To more selectively evaluate the contribution of tactile feedback in guiding dexterous motor output, we designed a novel head-fixed tactile orienting task for the mouse. The assay was loosely modelled after a task developed for use in humans to distinguish the acuity of tactile orienting from tactile perception (Pruszynski, Flanagan et al. 2018). The task requires mice to use the glabrous pad of the hand to detect the orientation of grooves on a textured platform and use this tactile information to rotate a movable platform to a prescribed target angle. Throughout behavioral shaping, the task becomes progressively harder in a closed-loop fashion (i.e., introduction of a larger range of random starting orientations, smaller target zones, and longer hold periods). The complete design and code for this assay are available at www.github.com/azimlabsalk.

Tactile orientation cues were provided by a 3D printed (Form 2, Formlabs) pedestal lined with parallel ridges. The pedestal was 16 mm in diameter with 16 parallel and evenly spaced ridges at 1 mm intervals. As a control, in some trials the textured pedestal was replaced with a 3D printed smooth pedestal 16 mm in diameter. The base of the pedestal was connected to a motor/encoder unit (precious metal brushes EBCCL, 3.2W, Maxon; Encoder MR type M, 256 counts per turn, Maxon) that controls and records the orientation of the pedestal. The motor/encoder unit is passively mobile through a 180° range (−30° to 150°, with 0° neutral angle defined as perpendicular to the body axis of the mouse), enabling mice to easily turn the pedestal when the motor is not actively engaged. During the task, the orientation of the pedestal ridges was actively manipulated through a closed-loop feedback system using an AutoPID library written for a microcontroller board (Arduino Mega 2560, Arduino) controlled through a custom-made PCB board. The PID regulates the pedestal angle by modulating motor torque based on feedback from the rotary encoder, including parameters such as current angle and angular velocity. Task related commands such as target angle, window play of target, task duration, and hold duration were controlled by a GUI task controller developed with the MATLAB data acquisition toolbox through a data acquisition device (NI DAQ USB- 6002, National Instruments). Behavior was continuously monitored using an infrared USB web camera (OV2710, OmniVision). The entire behavioral system was placed in a sound attenuation box (ENV- 022MD-27, Med Associates Inc.), and training and experimental sessions were performed under red light.

#### Head-fixation

The tactile orientation task was carried out in head-fixed mice that were water restricted and maintained at ∼85% of their original body weight. The procedures for implanting the headpost and for post-operative care were based on previously established methods (Guo, Hires et al. 2014). Briefly, mice were anesthetized with isoflurane and placed into a stereotaxic frame, as described above. Custom-made headposts were lowered onto the skull using a stereotaxic manipulator designed to hold the headpost, centered 0.5 mm anterior from bregma along the midline, and secured with epoxy. After ∼5 days of recovery, water restriction was initiated (1 ml/day, weight monitored daily), and mice were acclimated to head-fixation according to previously described methods (Guo, Hires et al. 2014, Galinanes, Bonardi et al. 2018).

#### Task structure

The final task was structured such that the trial was initiated when the pedestal was oriented to a reset position of −30° (30° aft of the axis perpendicular to the orientation of the mouse). Immediately after the reset, a randomly chosen starting orientation (0° ± 25°) was set and the trial was initiated by rendering the motor passive, enabling the mouse to freely move the pedestal. In all cases, the behavioral apparatus forced animals to use their right hand to accomplish the task. To receive a reward (8 μl water drop from a lick spout), the animal was required to orient the pedestal ridges to a defined target angle of 60° ± 20° by turning the platform clockwise and maintaining position within the target window for 600 msec. If the animal was unsuccessful in completing the task within 7 sec, the trial was terminated and the pedestal was reset to −30° without reward. The task continued until the mouse either performed 105 successful trials or until a total of 160 trials was attempted in a given day. If an animal became sated with water or if it struggled within the fixation frame, the task was terminated for the day.

#### Training period

After 5 days of acclimating to head-fixation, training on the tactile orientation task was initiated. During the first week of training, task requirements were made easier by assigning a more restrictive starting window (0° ± 5°), a more lenient target window (60° ± 40°), and a shorter hold duration (100 msec) as reward criteria. After mice successfully learned to turn the pedestal under these lenient criteria, task difficulty was incrementally increased towards the final task structure described above. To maximize training efficiency, reward criteria were constantly adjusted via a computer controlled interface based on performance. For example, if the success rate over a given 10 trials was > 80%, the reward criteria for the next 10 trials was incrementally adjusted to the next difficulty. Conversely, if the success rate was < 70% over 10 trials, the reward criteria for the next 10 trials was eased. Mice were considered trained when they were able to obtain > 60% success with the most stringent criteria (target window = 60° ± 20°; random starting orientation = 0 ± 25°; hold duration = 600 ms), which typically occurred after 4-6 weeks of training. Mice unable to attain 60% success on the most stringent criteria were excluded from the study.

#### Optogenetic manipulation of cuneate inhibitory neurons

To evaluate the impact of activating local cuneate inhibitory circuits, AAV5-hSyn-DIO-oChIEF-Citrine was injected into the cuneate region of adult *VGAT-Cre* mice, as described above. For control experiments, AAV5-CMV-EGFP was injected into the cuneate region of adult wild-type mice, as described above. Animals were left for at least 3 weeks before undergoing an additional surgery to implant optical fibers targeting the cuneate, as described above. Animals were returned to their home cage and allowed at least 1 week to recover before training was initiated. After mice were trained to perform the task, data acquisition trials were initiated. During the acquisition period, mice underwent 40 practice trials and then moved on to test trials in which they received photostimulation throughout the duration of the trial (473 nm laser, 10 Hz, 50 msec pulse width, ∼13 mW at patch cable interface). Photostimulation trials were randomly selected and occurred at a probability of 30-40%.

#### Data analysis

Data collected from each animal, including angular trajectories and time to complete trials, were concatenated for further analysis in MATLAB. The success rate was computed as the number of rewarded trials/total number of trials. To assess the efficiency of task execution specifically on successful trials, we excluded failures and computed the cumulative distribution functions (CDF) of the time to complete each trial. To compare CDF data between mice, we randomly selected a fixed number of samples from each animal (determined by the animal with the lowest number of successful trials).

### Statistics and data collection

Mice from each litter were randomly allocated to different groups for the electrophysiological and behavioral experiments. Group sizes were not pre-determined, but sample sizes are comparable to those commonly used for similar experiments, and were selected such that appropriate statistical tests could be used. Data analysis for extracellular recording and string pulling behavior were blinded. In the tactile orienting assay, data collection and analysis were automated, minimizing any potential influence by the experimenter. Animals were only excluded from analysis if it was found that viral targeting or fiber placement were off target, if mice became unhealthy, or if they did not achieve baseline level of behavioral performance before perturbation. Results are shown as box-and whisker plots indicating the median, 25^th^ and 75^th^ percentiles, and range, unless otherwise indicated. Normality was assessed using the Shapiro-Wilk test or the Kolmogorov-Smirnov test, and non-parametric tests were used for non-normally distributed data. All parametric and non-parametric tests used, as well as any multiple comparisons tests and corrections are indicated in the figure legends, as are *n* values for each experiment. All statistical comparisons were two-tailed, when relevant. P < 0.05 was considered significant, * indicates *P* < 0.05, ** < 0.01, *** < 0.001, **** < 0.001, and all significant P values are indicated in figure legends. Statistical analysis was performed in MATLAB or Prism (version 8.4.3, GraphPad).

## References

Adams, R. A., S. Shipp and K. J. Friston (2013). “Predictions not commands: active inference in the motor system.” Brain Struct Funct 218(3): 611–643.

Aguilar, J., C. Rivadulla, C. Soto and A. Canedo (2003). “New corticocuneate cellular mechanisms underlying the modulation of cutaneous ascending transmission in anesthetized cats.” J Neurophysiol 89(6): 3328–3339.

Andersen, P., J. C. Eccles, T. Oshima and R. F. Schmidt (1964). “Mechanisms of Synaptic Transmission in the Cuneate Nucleus.” J Neurophysiol 27: 1096–1116.

Andersen, P., J. C. Eccles and R. F. Schmidt (1962). “Presynaptic inhibition in the cuneate nucleus.” Nature 194: 741–743.

Azim, E. and K. Seki (2019). “Gain control in the sensorimotor system.” Curr Opin Physiol 8: 177–187.

Ballermann, M., J. McKenna and I. Q. Whishaw (2001). “A grasp-related deficit in tactile discrimination following dorsal column lesion in the rat.” Brain Res Bull 54(2): 237–242.

Bengtsson, F., R. Brasselet, R. S. Johansson, A. Arleo and H. Jorntell (2013). “Integration of sensory quanta in cuneate nucleus neurons in vivo.” PLoS One 8(2): e56630.

Berkley, K. J., R. J. Budell, A. Blomqvist and M. Bull (1986). “Output systems of the dorsal column nuclei in the cat.” Brain Res 396(3): 199–225.

Blackwell, A. A., M. T. Banovetz, Qandeel, I. Q. Whishaw and D. G. Wallace (2018). “The structure of arm and hand movements in a spontaneous and food rewarded on-line string-pulling task by the mouse.” Behav Brain Res 345: 49–58.

Blakemore, S. J., C. D. Frith and D. M. Wolpert (1999). “Spatio-temporal prediction modulates the perception of self-produced stimuli.” J Cogn Neurosci 11(5): 551–559.

Canedo, A. (1997). “Primary motor cortex influences on the descending and ascending systems.” Prog Neurobiol 51(3): 287–335.

Canedo, A., J. Marino and J. Aguilar (2000). “Lemniscal recurrent and transcortical influences on cuneate neurons.” Neuroscience 97(2): 317–334.

Chapin, J. K. and D. J. Woodward (1981). “Modulation of sensory responsiveness of single somatosensory cortical cells during movement and arousal behaviors.” Exp Neurol 72(1): 164–178.

Chapman, C. E. (1994). “Active versus passive touch: factors influencing the transmission of somatosensory signals to primary somatosensory cortex.” Can J Physiol Pharmacol 72(5): 558–570.

Cheema, S., B. L. Whitsel and A. Rustioni (1983). “The corticocuneate pathway in the cat: relations among terminal distribution patterns, cytoarchitecture, and single neuron functional properties.” Somatosens Res 1(2): 169–205.

Cole, J. D. and G. Gordon (1992). “Corticofugal actions on lemniscal neurons of the cuneate, gracile and lateral cervical nuclei of the cat.” Exp Brain Res 90(2): 384–392.

Confais, J., G. Kim, S. Tomatsu, T. Takei and K. Seki (2017). “Nerve-Specific Input Modulation to Spinal Neurons during a Motor Task in the Monkey.” J Neurosci 37(10): 2612–2626.

Coulter, J. D. (1974). “Sensory transmission through lemniscal pathway during voluntary movement in the cat.” J Neurophysiol 37(5): 831–845.

Delaurier, A., N. Burton, M. Bennett, R. Baldock, D. Davidson, T. J. Mohun and M. P. Logan (2008). “The Mouse Limb Anatomy Atlas: an interactive 3D tool for studying embryonic limb patterning.” BMC Dev Biol 8: 83.

Fink, A. J., K. R. Croce, Z. J. Huang, L. F. Abbott, T. M. Jessell and E. Azim (2014). “Presynaptic inhibition of spinal sensory feedback ensures smooth movement.” Nature 509(7498): 43–48.

Ghez, C. and M. Pisa (1972). “Inhibition of afferent transmission in cuneate nucleus during voluntary movement in the cat.” Brain Res 40(1): 145–155.

Gilbert, C. D. and M. Sigman (2007). “Brain states: top-down influences in sensory processing.” Neuron 54(5): 677–696.

Glendinning, D. S., B. Y. Cooper, C. J. Vierck, Jr. and C. M. Leonard (1992). “Altered precision grasping in stumptail macaques after fasciculus cuneatus lesions.” Somatosens Mot Res 9(1): 61–73.

Hantman, A. W. and T. M. Jessell (2010). “Clarke’s column neurons as the focus of a corticospinal corollary circuit.” Nat Neurosci 13(10): 1233–1239.

Hippenmeyer, S., E. Vrieseling, M. Sigrist, T. Portmann, C. Laengle, D. R. Ladle and S. Arber (2005). “A developmental switch in the response of DRG neurons to ETS transcription factor signaling.” PLoS Biol 3(5): e159.

Jabbur, S. J. and A. L. Towe (1960). “Effect of pyramidal tract activity on dorsal column nuclei.” Science 132(3426): 547–548.

Johansson, R. S. and J. R. Flanagan (2009). “Coding and use of tactile signals from the fingertips in object manipulation tasks.” Nat Rev Neurosci 10(5): 345–359.

Jorntell, H., F. Bengtsson, P. Geborek, A. Spanne, A. V. Terekhov and V. Hayward (2014). “Segregation of tactile input features in neurons of the cuneate nucleus.” Neuron 83(6): 1444–1452.

Keller, G. B. and T. D. Mrsic-Flogel (2018). “Predictive Processing: A Canonical Cortical Computation.” Neuron 100(2): 424–435.

Kuypers, H. G. and J. D. Tuerk (1964). “The Distribution of the Cortical Fibres within the Nuclei Cuneatus and Gracilis in the Cat.” J Anat 98: 143–162.

Lee, S., G. E. Carvell and D. J. Simons (2008). “Motor modulation of afferent somatosensory circuits.” Nat Neurosci 11(12): 1430–1438.

Liu, Y., A. Latremoliere, X. Li, Z. Zhang, M. Chen, X. Wang, C. Fang, J. Zhu, C. Alexandre, Z. Gao, B. Chen, X. Ding, J. Y. Zhou, Y. Zhang, C. Chen, K. H. Wang, C. J. Woolf and Z. He (2018). “Touch and tactile neuropathic pain sensitivity are set by corticospinal projections.” Nature 561(7724): 547–550.

Loutit, A. J., R. M. Vickery and J. R. Potas (2020). “Functional organization and connectivity of the dorsal column nuclei complex reveals a sensorimotor integration and distribution hub.” J Comp Neurol.

Lue, J. H., Y. F. Jiang-Shieh, J. Y. Shieh, E. A. Ling and C. Y. Wen (1997). “Multiple inputs of GABA-immunoreactive neurons in the cuneate nucleus of the rat.” Neurosci Res 27(2): 123–132.

Mathis, A., P. Mamidanna, K. M. Cury, T. Abe, V. N. Murthy, M. W. Mathis and M. Bethge (2018). “DeepLabCut: markerless pose estimation of user-defined body parts with deep learning.” Nat Neurosci 21(9): 1281–1289.

McComas, A. J. (2016). “Hypothesis: Hughlings Jackson and presynaptic inhibition: is there a big picture?” J Neurophysiol 116(1): 41–50.

Milne, R. J., A. M. Aniss, N. E. Kay and S. C. Gandevia (1988). “Reduction in perceived intensity of cutaneous stimuli during movement: a quantitative study.” Exp Brain Res 70(3): 569–576.

Miyamichi, K., Y. Shlomai-Fuchs, M. Shu, B. C. Weissbourd, L. Luo and A. Mizrahi (2013). “Dissecting local circuits: parvalbumin interneurons underlie broad feedback control of olfactory bulb output.” Neuron 80(5): 1232–1245.

Mountcastle, V. B. (2005). The Sensory Hand : Neural Mechanisms of Somatic Sensation. Cambridge, Mass., Harvard University Press.

Nath, T., A. Mathis, A. C. Chen, A. Patel, M. Bethge and M. W. Mathis (2019). “Using DeepLabCut for 3D markerless pose estimation across species and behaviors.” Nat Protoc 14(7): 2152–2176.

Niu, J., L. Ding, J. J. Li, H. Kim, J. Liu, H. Li, A. Moberly, T. C. Badea, I. D. Duncan, Y. J. Son, S. S. Scherer and W. Luo (2013). “Modality-based organization of ascending somatosensory axons in the direct dorsal column pathway.” J Neurosci 33(45): 17691–17709.

Popratiloff, A., J. G. Valtschanoff, A. Rustioni and R. J. Weinberg (1996). “Colocalization of GABA and glycine in the rat dorsal column nuclei.” Brain Res 706(2): 308–312.

Pruszynski, J. A., J. R. Flanagan and R. S. Johansson (2018). “Fast and accurate edge orientation processing during object manipulation.” Elife 7.

Rustioni, A. and N. L. Hayes (1981). “Corticospinal tract collaterals to the dorsal column nuclei of cats. An anatomical single and double retrograde tracer study.” Exp Brain Res 43(3-4): 237–245.

Rustioni, A., D. E. Schmechel, S. Cheema and D. Fitzpatrick (1984). “Glutamic acid decarboxylase-containing neurons in the dorsal column nuclei of the cat.” Somatosens Res 1(4): 329–357.

Schneider, D. M., J. Sundararajan and R. Mooney (2018). “A cortical filter that learns to suppress the acoustic consequences of movement.” Nature 561(7723): 391–395.

Scott, S. H. (2016). “A Functional Taxonomy of Bottom-Up Sensory Feedback Processing for Motor Actions.” Trends Neurosci 39(8): 512–526.

Shadmehr, R., M. A. Smith and J. W. Krakauer (2010). “Error correction, sensory prediction, and adaptation in motor control.” Annu Rev Neurosci 33: 89–108.

Shin, H. C. and J. K. Chapin (1989). “Mapping the effects of motor cortex stimulation on single neurons in the dorsal column nuclei in the rat: direct responses and afferent modulation.” Brain Res Bull 22(2): 245–252.

Sillito, A. M., H. E. Jones, G. L. Gerstein and D. C. West (1994). “Feature-linked synchronization of thalamic relay cell firing induced by feedback from the visual cortex.” Nature 369(6480): 479–482.

Soto, C., J. Aguilar, F. Martin-Cora, C. Rivadulla and A. Canedo (2004). “Intracuneate mechanisms underlying primary afferent cutaneous processing in anaesthetized cats.” Eur J Neurosci 19(11): 3006–3016.

Suresh, A. K., J. E. Winberry, C. Versteeg, R. Chowdhury, T. Tomlinson, J. M. Rosenow, L. E. Miller and S. J. Bensmaia (2017). “Methodological considerations for a chronic neural interface with the cuneate nucleus of macaques.” J Neurophysiol 118(6): 3271–3281.

Towe, A. L. and S. J. Jabbur (1961). “Cortical inhibition of neurons in dorsal column nuclei of cat.” J Neurophysiol 24: 488–498.

Wall, P. D. (1970). “The sensory and motor role of impulses travelling in the dorsal columns towards cerebral cortex.” Brain 93(3): 505–524.

Witham, C. L. and S. N. Baker (2011). “Modulation and transmission of peripheral inputs in monkey cuneate and external cuneate nuclei.” J Neurophysiol 106(5): 2764–2775.

Zhou, X., L. Wang, H. Hasegawa, P. Amin, B. X. Han, S. Kaneko, Y. He and F. Wang (2010). “Deletion of PIK3C3/Vps34 in sensory neurons causes rapid neurodegeneration by disrupting the endosomal but not the autophagic pathway.” Proc Natl Acad Sci U S A 107(20): 9424–9429.

