## Supplementary Figures for "Modulation of tactile feedback for the execution of dexterous movement"

Eiman Azim[1*]

[1]Molecular Neurobiology Laboratory, Salk Institute for Biological Studies; La Jolla, CA, USA.

[†]These authors contributed equally to this work

### Supplementary Figures

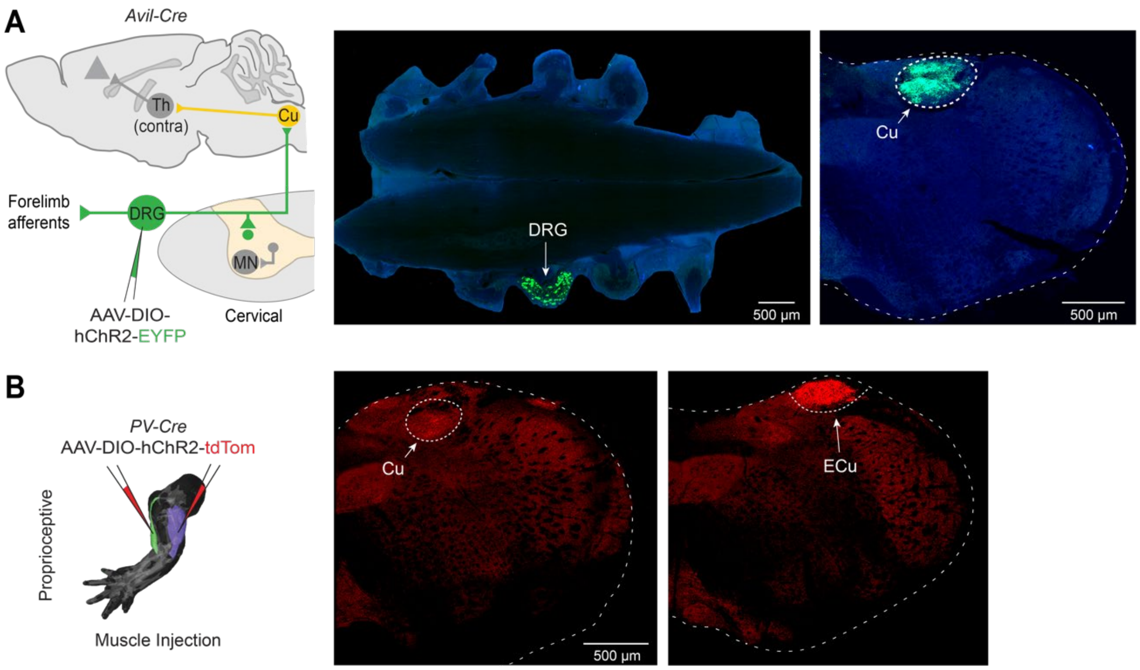


**Supplementary Figure 1. Termination of ascending forelimb sensory afferents in the dorsal column nuclei.**

**(A)** Labeling of sensory afferents through injection of AAV-DIO-hChR2-EYFP or AAV-CMV-EGFP (not shown) into a single dorsal root ganglion (DRG, C6 or C7) of *Avil-Cre* mice reveals dense projections to the ipsilateral Cu (4 mice). The *Avil-Cre* mouse line drives expression selectively and broadly across sensory neuron classes (Zhou, Wang et al. 2010). **(B)** Selective labeling of proprioceptive neurons though injection of AAV-DIO-hChR2-tdTom or AAV-DIO-EGFP (not shown) into forelimb muscles (biceps and triceps) of *PV-Cre* mice reveals minimal targeting of Cu, but extensive targeting of the ipsilateral ECu and some projections to other regions of the cuneate nucleus (not shown) (5 mice). The *Pv-Cre* mouse line drives sensory neuron expression that is largely specific to proprioceptors (Hippenmeyer, Vrieseling et al. 2005). Limb muscle image from (Delaurier, Burton et al. 2008).

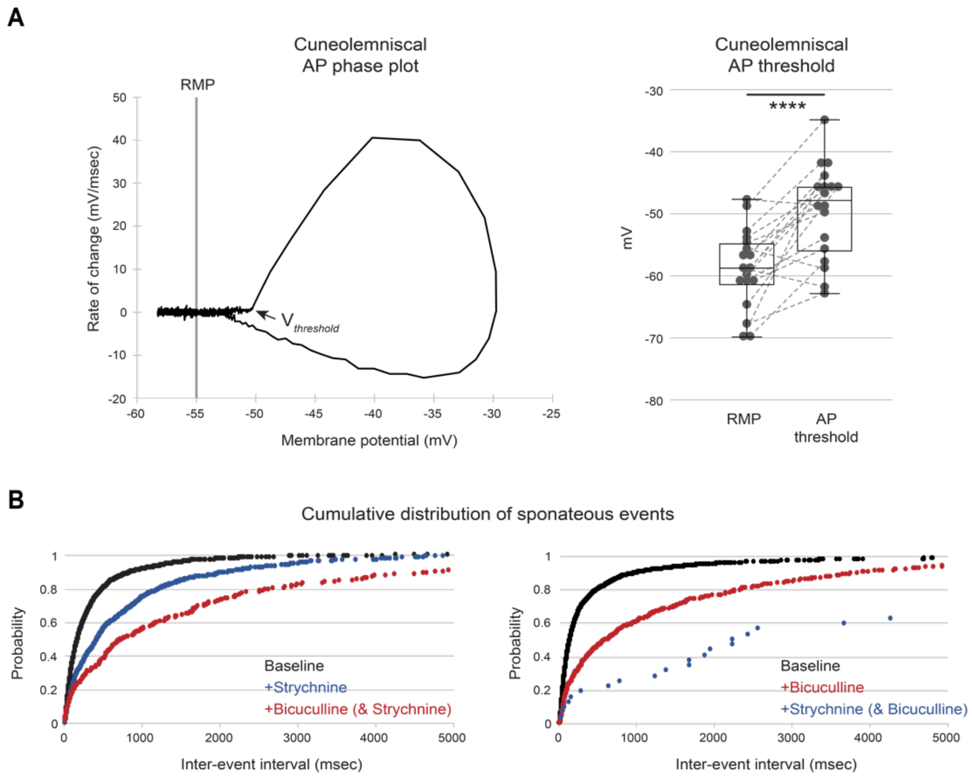


**Supplementary Figure 2. Cuneolemniscal neuron action potential threshold and spontaneous inhibitory input.**
**(A)** Whole-cell slice recordings from labeled cuneolemniscal (CL) neurons (as in **Fig. 1C**). Example phase plot (left) showing the rate of change of membrane potential versus membrane potential. The voltage threshold ($V_{threshold}$) for firing an action potential (AP) was calculated from the first sweep of current injection that produced an AP. Resting membrane potential (RMP, -59.06 mV ± 1.50 mV SEM) is slightly less than AP threshold (-49.67 mV ± 1.78 mV SEM) (right, 18 neurons from 10 mice; ****P < 0.0001; paired t test; also see **Fig. 1D**). **(B)** Cumulative probability distribution of the inter-event intervals of spontaneous postsynaptic currents recorded from CL neurons at baseline (black), and followed by sequential bath application of strychnine (blue) then bicuculline (red) (left, 7 neurons in 4 mice), or bicuculline then strychnine (right, 8 neurons in 4 mice).

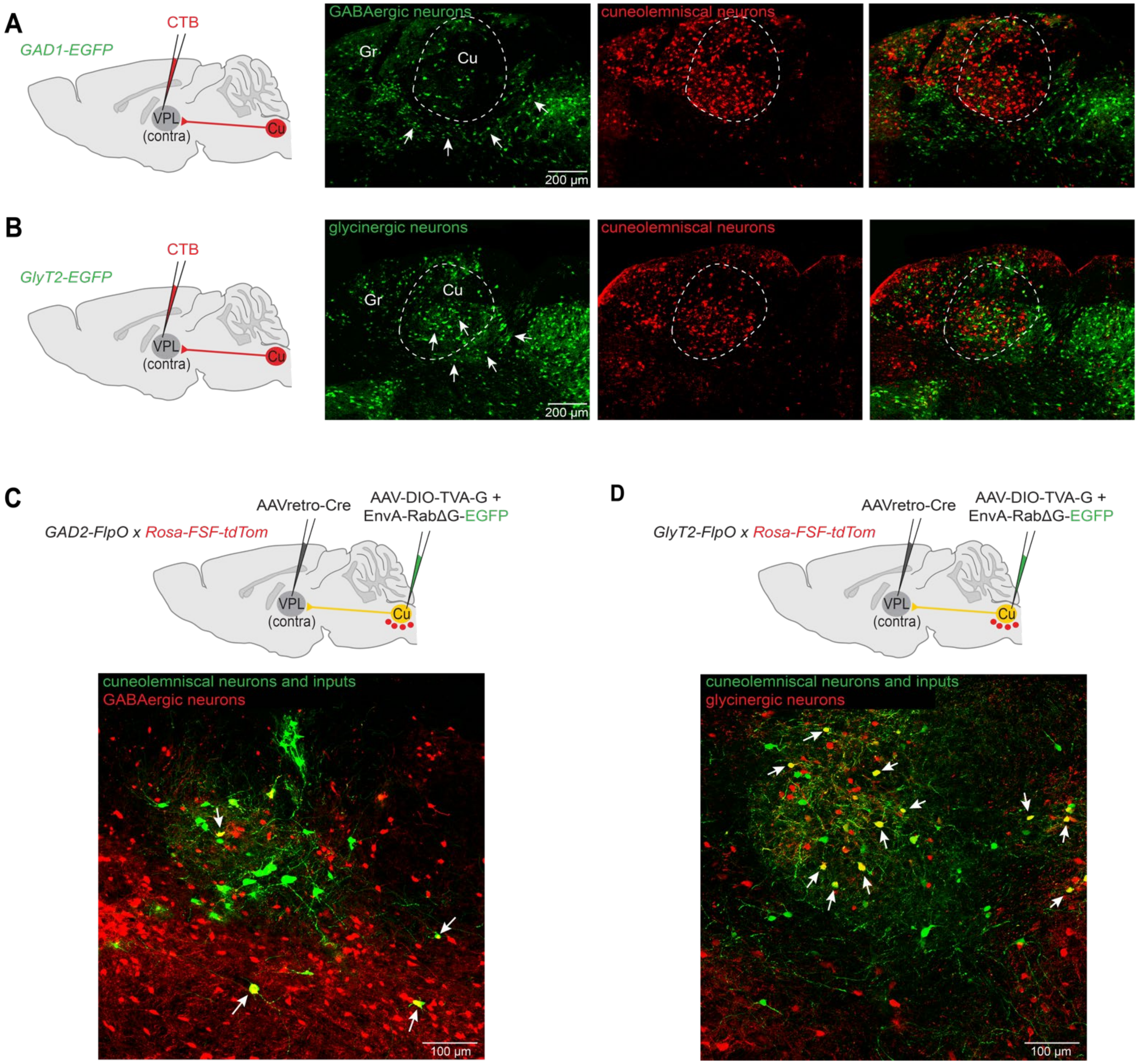


**Supplementary Figure 3. GABAergic and glycinergic neurons are differentially distributed within the cuneate region and directly target cuneolemniscal neurons.**

**(A)** *GAD1-EGFP* mice were used to label GABAergic neurons (green), and cuneolemniscal (CL) neurons (red) were traced by injecting either CTB or Fluorogold (not shown) into contralateral VPL thalamus (3 mice). Few GABAergic neurons are localized within the Cu core region, where CL neurons are found, but are instead located primarily within the ventral shell of the Cu (arrows). In no cases were GABAergic neurons co-labeled with CTB or Fluorogold. Overlay shown on right. Gr, gracile nucleus. **(B)** *GlyT2-EGFP* mice were used to label glycinergic neurons (green), and CL neurons (red) were traced by injecting either CTB or Fluorogold (not shown) into contralateral VPL thalamus (3 mice). Many glycinergic neurons can be found within the Cu core region as well as in the ventral shell region (arrows). In no cases were glycinergic neurons co-labeled with CTB or Fluorogold. Overlay shown on right. It is possible that some of these inhibitory neurons are positive for both glycine and GABA (Popratiloff, Valtschanoff et al.

1996). **(C)** Monosynaptic retrograde rabies tracing from CL neurons through injection of AAVretro-Cre into VPL thalamus and AAV-DIO-TVA-G into Cu, followed 3-4 weeks later by injection of EnvA-pseudotyped RabΔG-EGFP into Cu (2 mice). GABAergic cells (red) were labeled by crossing *GAD2-FlpO* mice with a *Rosa-FSF-tdTom* line. Yellow neurons (arrows) indicate GABAergic neurons, located largely in the Cu ventral shell, that provide monosynaptic inputs to CL neurons. Note that use of a TC66T strain of helper virus, containing a mutated, low-affinity form of the TVA receptor, limits nonspecific rabies transduction due to low levels of leaky TVA expression (Miyamichi, Shlomai-Fuchs et al. 2013). **(D)** Monosynaptic retrograde rabies tracing from CL neurons (as in (C); 2 mice). Glycinergic cells (red) were labeled by crossing *GlyT2-FlpO* mice with a *Rosa-FSF-tdTom* line. Yellow neurons (arrows) indicate glycinergic neurons located in the Cu core and ventral shell that provide monosynaptic inputs to CL neurons.

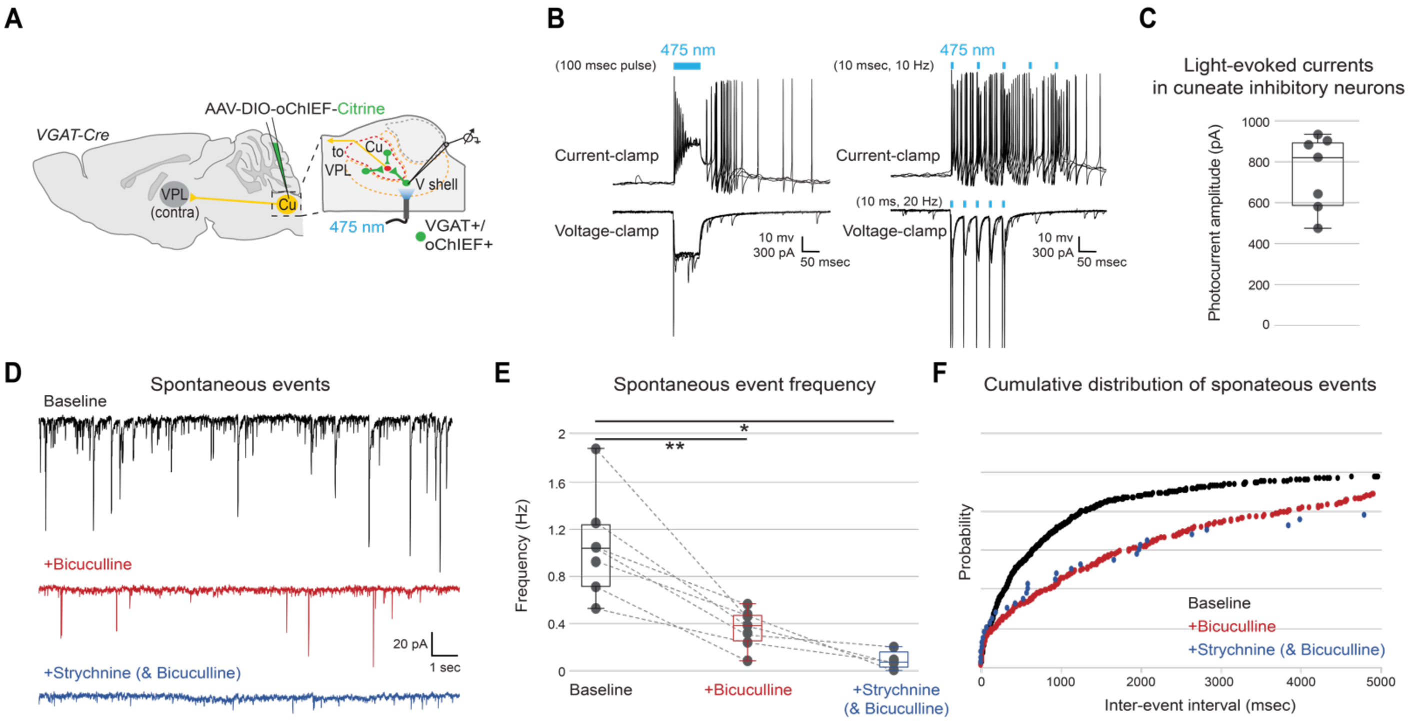


**Supplementary Figure 4. Optogenetic activation and spontaneous inhibition in cuneate inhibitory neurons.**
**(A)** *In vitro* slice recording from Cu inhibitory neurons optogenetically activated following injection of AAV-DIO-oChIEF-Citrine in the Cu core and ventral shell (V shell) regions of *VGAT-Cre* mice. Inhibitory neurons were targeted for recording by Citrine expression. **(B)** Example whole-cell recording from a labeled inhibitory neuron in current clamp (top) and voltage clamp (bottom), with a continuous 100 msec light pulse (left), or five 10 msec light pulses at 10 or 20 Hz (right). Light-evoked responses are observed under all conditions. **(C)** Quantification of light-evoked currents (100 msec light pulse, 7 neurons in 6 mice). **(D)** Example whole-cell recording from a Citrine-positive inhibitory neuron held at -70mV showing spontaneous events at baseline (black). Bath application of bicuculline (red) followed by strychnine (blue) progressively eliminates most spontaneous inhibitory post-synaptic currents (IPSCs). Pipette was filled with high chloride solution, causing IPSCs to appear as inward currents. **(E)** Sequential application of bicuculline and strychnine cumulatively decreases the frequency of spontaneous IPSCs in Cu inhibitory neurons (7 neurons in 5 mice, 4 neurons for strychnine; ***P* = 0.0063, **P* = 0.0147; One-way mixed-effects model with Geisser-Greenhouse correction and Tukey's multiple comparisons test). **(F)** Cumulative probability distribution of the inter-event intervals of spontaneous postsynaptic currents recorded from Cu inhibitory neurons at baseline (black), and followed by sequential bath application of bicuculline (red) then strychnine (blue) (7 neurons in 5 mice, 4 neurons for strychnine). Subsequent application of strychnine minimally alters event probability, suggesting the majority of IPSCs are $GABA_A$ receptor mediated.

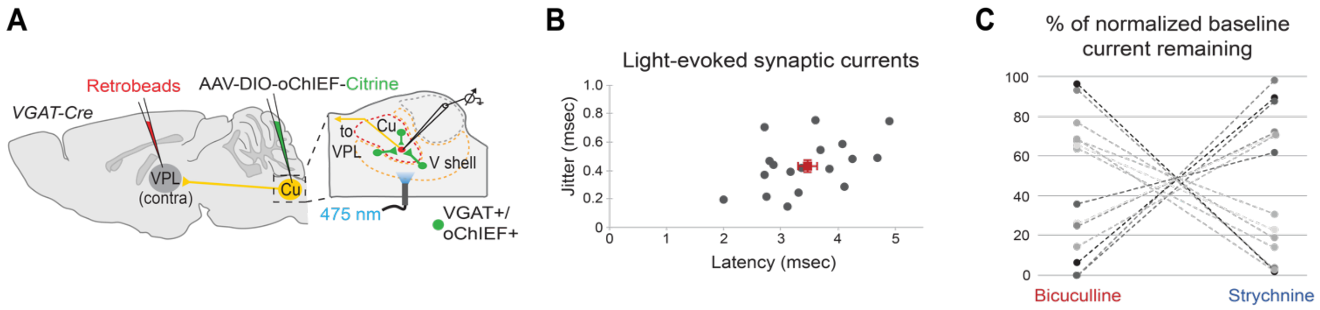


**Supplementary Figure 5. The relative ratio of GABAergic and glycinergic monosynaptic inhibition of cuneolemniscal neurons varies across cells.**
**(A)** *In vitro* slice recording from CL neurons retrogradely labeled by retrobead injection into contralateral VPL thalamus. Cu inhibitory neurons were optogenetically activated following injection of AAV-DIO-oChIEF-Citrine in the Cu of *VGAT-Cre* mice (as in **Fig. 1H**). **(B)** Onset kinetics of light-evoked IPSCs in CL neurons. Plot shows short latency and low jitter (standard deviation of trial-to-trial event latencies) of light-evoked IPSCs (error bars indicate SEM, 18 neurons in 8 mice). **(C)** Sequential application of bicuculline then strychnine or strychnine then bicuculline shows cumulative reduction in the amplitude of light-evoked IPSCs with an approximately equal mix of GABAergic and glycinergic components (see **Fig. 1I**). The ratio of GABAergic and glycinergic inhibition onto CL neurons varies across cells (14 neurons in 7 mice). In neurons where GABAergic inhibition is low, glycinergic inhibition is high, and vice versa. The order of strychnine and bicuculline application did not alter the combined effect.

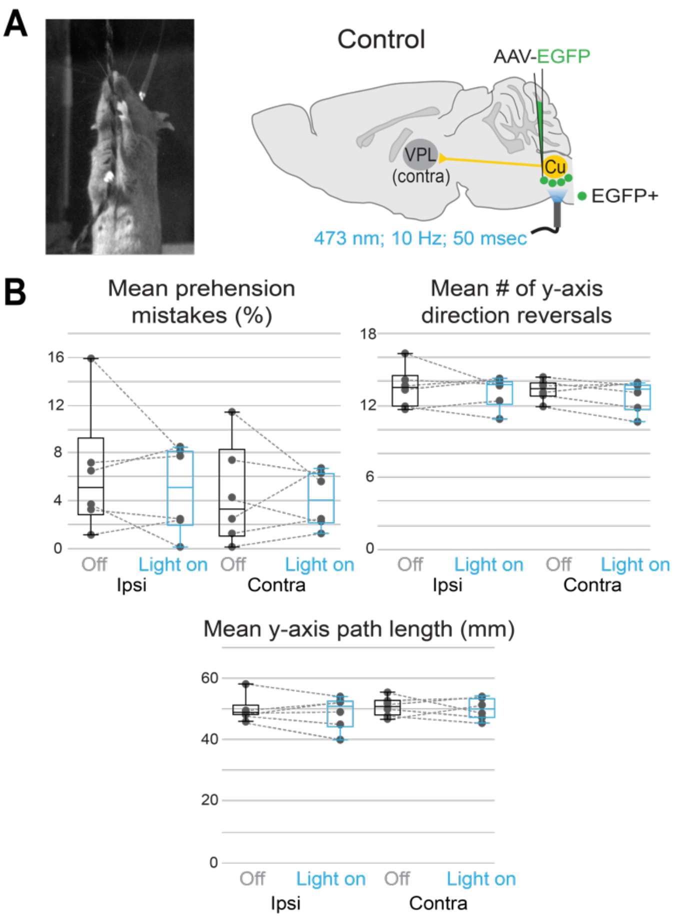


**Supplementary Figure 6. Photostimulation in control mice does not affect string pull performance or kinematics.**
**(A)** String pulling task (left). As a control, wild-type mice received unilateral injection of AAV-EGFP into the Cu region followed by implantation of an optical fiber (also see **Fig. 3A-D**). **(B)** Photostimulation has no effect on ipsilateral or contralateral prehension mistakes (top left; % of grasp attempts in which an error was made across trials, see Materials and Methods), the mean number of vertical (y) direction reversals (top right; mean number of direction reversals per trial), or the mean y path length traversed by either hand between direction reversals (bottom; mean absolute distance between the peak and trough of a given path segment across trials) (6 mice; Two-way repeated measures ANOVA with Sidak multiple comparisons test).

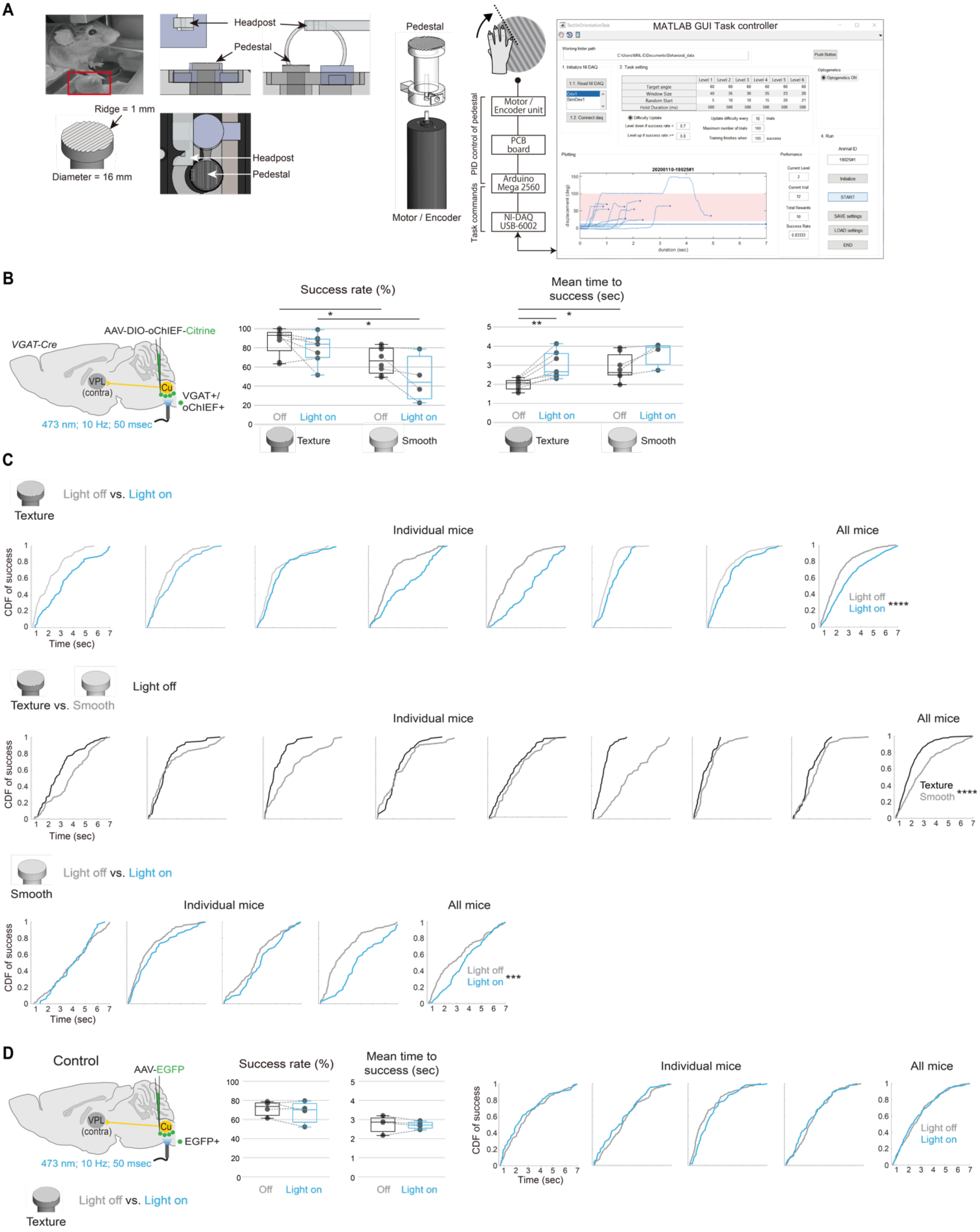
A
Headpost
Pedestal
Ridge = 1 mm
Diameter = 16 mm
Headpost
Pedestal
Pedestal
Motor / Encoder
PID control of pedestal
Task commands
Motor / Encoder unit
PCB board
Arduino Mega 2560
NI-DAQ USB-6002
MATLAB GUI Task controller
B
VGAT-Cre
AAV-DIO-oChIEF-Citrine
VPL (contra)
Cu
VGAT+/ oChIEF+
473 nm; 10 Hz; 50 msec
Success rate (%)
Mean time to success (sec)
Off
Light on
Texture
Smooth
C
Light off vs. Light on
Texture
Individual mice
All mice
CDF of success
Time (sec)
Light off
Light on
****
Texture vs. Smooth
Light off
Texture
Smooth
****
Light off vs. Light on
Smooth
***
D
Control
AAV-EGFP
VPL (contra)
Cu
EGFP+
473 nm; 10 Hz; 50 msec
Light off vs. Light on
Texture
Success rate (%)
Mean time to success (sec)
Off
Light on
Individual mice
All mice
CDF of success
Time (sec)
Light off
Light on

**Supplementary Figure 7. Aberrant activation of cuneate inhibitory circuits disrupts performance of a tactile orientation task.**

**(A)** Schematic of head-fixed, tactile orienting assay. A pedestal (16 mm diameter) with parallel orientation ridges (1 mm interval, evenly spaced) is placed under the right hand. A headpost is used to restrain the mouse. The pedestal is connected to a motor/rotary encoder assembly. A GUI task controller provides PID control of the pedestal, delivers task commands, and records performance (see Materials and Methods). **(B)** Ipsilateral Cu inhibitory neurons were targeted for optogenetic activation (left, as in **Fig. 4**). During photoactivation (blue), the overall success rate (middle) is unaffected in mice using either a textured (7 mice paired) or a smooth surface (8 mice with light off, 4 mice with light on; 4 paired). However, success rate is reduced when mice switch from a textured to a smooth surface with the light off (8 mice paired; **P* = 0.0355) or light on (7 mice with texture, 4 mice with smooth; 2 paired; **P* = 0.0245). The mean elapsed time to achieve success (right) increases during photoactivation for mice using a textured surface (7 mice paired; (***P* = 0.0047), and for mice that switch from a textured to a smooth surface with the light off (8 mice paired; **P* = 0.0170). (Two-way mixed-effects model with Geisser-Greenhouse correction and Sidak multiple comparisons test; also see **Fig. 4F-H**). Portions of this data are also presented in **Fig. 4G**. **(C)** The cumulative distribution function (CDF) for only the successful trials under each condition for individual mice and across all mice (right). A rightward shift of the CDF is seen during photoactivation in mice using a textured surface (7 mice; *****P* < 0.0001), when mice switch from a textured to a smooth surface with the light off (8 mice; *****P* < 0.0001), and when mice switch from a textured to a smooth surface with the light on (4 mice; ****P* < 0.0007). (Kolmogorov-Smirnov test; also see **Fig. 4I**). **(D)** As a control, wild-type mice received unilateral injection of AAV-EGFP into the Cu region followed by implantation of an optical fiber. Photostimulation in mice using a textured surface did not affect the task success rate (4 mice; paired t test), the mean elapsed time to achieve success (paired t test), or the CDF of successful trials (Kolmogorov-Smirnov test).

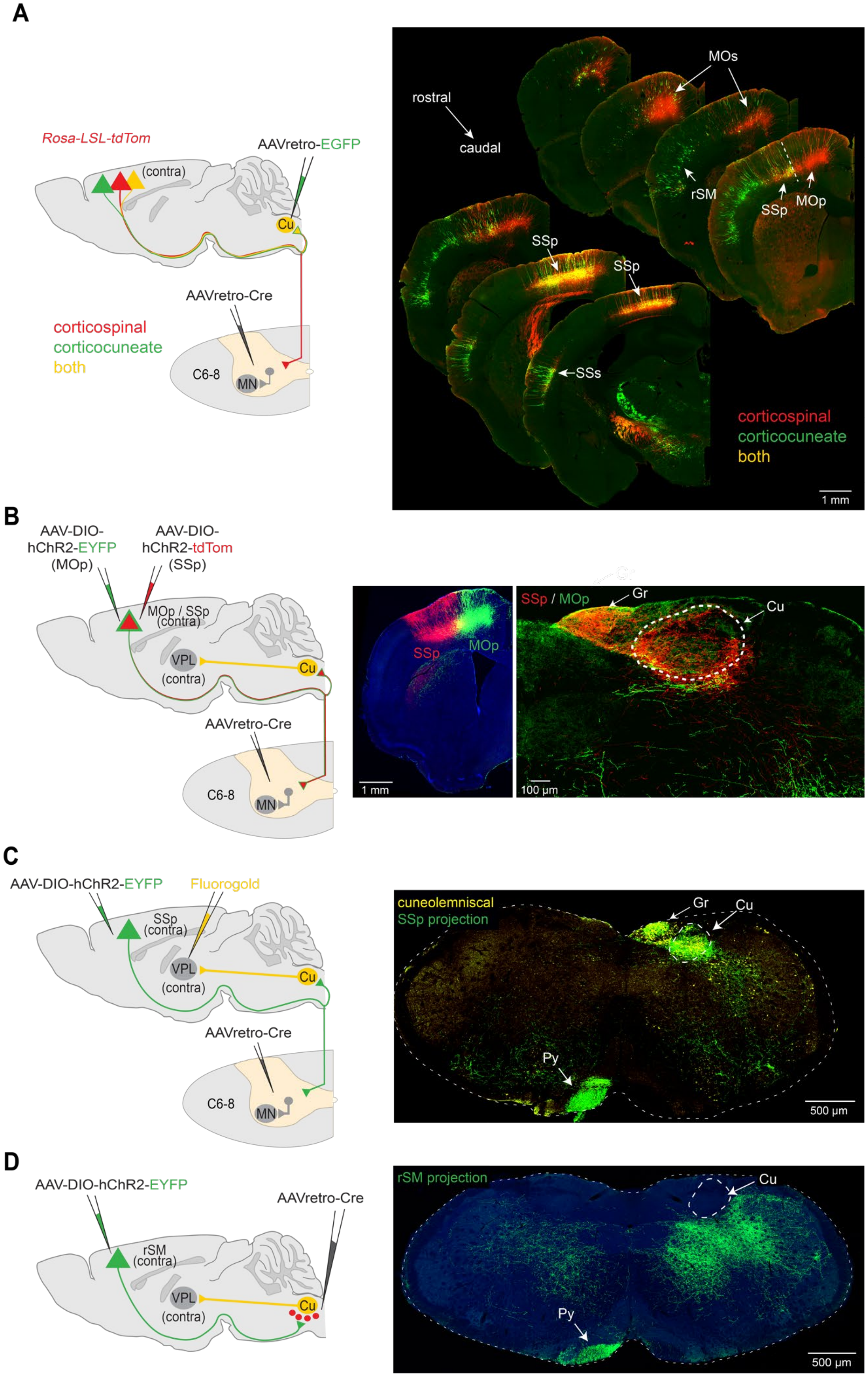
A
Rosa-LSL-tdTom
AAVretro-EGFP
(contra)
Cu
AAVretro-Cre
corticospinal
corticocuneate
both
C6-8
MN
MOs
rostral
caudal
rSM
SSp
MOp
SSs
1 mm
B
AAV-DIO-hChR2-EYFP (MOp)
AAV-DIO-hChR2-tdTom (SSp)
MOp / SSp (contra)
VPL
(contra)
Cu
AAVretro-Cre
C6-8
MN
SSp
MOp
1 mm
SSp / MOp
Gr
Cu
100 µm
C
AAV-DIO-hChR2-EYFP
Fluorogold
SSp (contra)
VPL
(contra)
Cu
AAVretro-Cre
C6-8
MN
cuneolemniscal
SSp projection
Gr
Cu
Py
500 µm
D
AAV-DIO-hChR2-EYFP
AAVretro-Cre
rSM (contra)
VPL
(contra)
Cu
rSM projection
Cu
Py
500 µm

**Supplementary Figure 8. Identifying corticofugal and corticospinal inputs to the cuneate region.** **(A)** Approach to broadly identify corticofugal and corticospinal inputs to the Cu region (left). AAVretro-Cre was injected into C6-C8 spinal segments in *Rosa-LSL-tdTom* mice, labeling corticospinal neurons (red). AAVretro-GFP was injected into the Cu to label descending corticocuneate projections (green). Corticospinal neurons that also target Cu are dual labeled (yellow). Corticospinal neurons (red) can be seen throughout contralateral primary (SSp) and supplemental (SSs) somatosensory cortices, as well as primary (MOp) and secondary motor cortices (MOs). Retrograde labeling from the Cu region showed considerable overlap with corticospinal projections (yellow), especially throughout SSp and SSs. Corticocuneate neurons (green) were found in contralateral rostral sensorimotor cortex (rSM), where very few corticospinal projection neurons are located. Corticocuneate neurons were rarely found within MOp or MOs. (3 mice). **(B)** Approach to distinguish projections from SSp and MOp cortices (left). To label corticospinal projections, AAVretro-Cre was injected into C6-C8 spinal segments, and AAV-DIO-hChR2-EYFP was injected into contralateral MOp and AAV-DIO-hChR2-tdTom (or AAV-DIO-hChR2-mCherry, data not shown) was injected into contralateral SSp. Fluorophore expression was segregated to the two cortical regions (middle). The core regions of contralateral Cu and Gr (right) are heavily innervated by SSp (red), while only sparse innervation from MOp (green) can be found, mostly in the ventral shell region. In some mice the viruses were switched, with no change in results. (4 mice). **(C)** Approach to identify projections to the Cu region that specifically arise from corticospinal neurons (left). AAVretro-Cre was injected into C6-C8 spinal segments, and AAV-DIO-hChR2-EYFP was injected into contralateral SSp. CL neurons were retrogradely labeled by Fluorogold injection into contralateral VPL thalamus. Corticospinal neurons in SSp descend through the ipsilateral pyramidal tract (Py), decussate, and send collaterals that densely target the core regions of contralateral Cu and Gr, where CL neurons are located (also see **Fig. 5B**). (6 mice). **(D)** Approach to identify projections to the Cu region that specifically arise from rSM (left). AAVretro-Cre was injected into the Cu region and AAV-DIO-hChR2-EYFP was injected into contralateral rSM. rSM projections also descend through the ipsilateral Py and decussate. However, unlike SSp, rSM projections completely avoid the core regions of Cu and instead target more ventral brainstem regions, including the Cu ventral shell region where inhibitory neurons that target CL neurons are located (also see **Fig. 5B**). (5 mice).

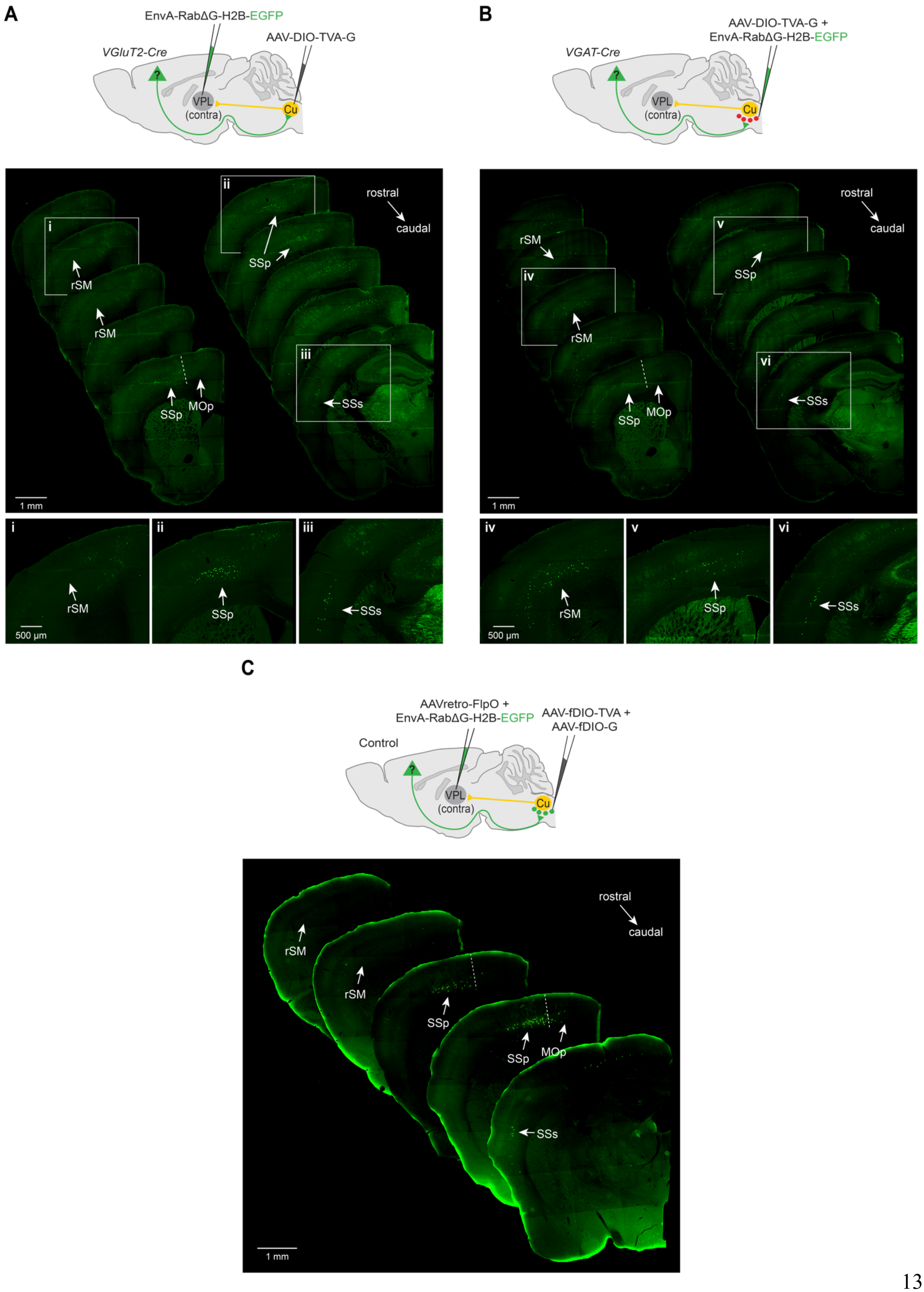
A
EnvA-RabΔG-H2B-EGFP
AAV-DIO-TVA-G
VGluT2-Cre
VPL
(contra)
Cu
rostral
caudal
i
ii
iii
rSM
SSp
MOp
SSs
1 mm
500 µm
B
AAV-DIO-TVA-G +
EnvA-RabΔG-H2B-EGFP
VGAT-Cre
iv
v
vi
C
AAVretro-FlpO +
EnvA-RabΔG-H2B-EGFP
AAV-fDIO-TVA +
AAV-fDIO-G
Control

**Supplementary Figure 9. Identifying brain-wide inputs onto cuneolemniscal neurons and cuneate inhibitory neurons.**

**(A)** A complementary monosynaptic rabies tracing strategy was used to validate the first approach (see **Fig. 5A**) and deliver nuclear localized fluorophore to enable automated quantification of labeled neurons across the brain. AAV-DIO-TVA-G was injected into Cu of *VGluT2-Cre* mice (avoiding transduction of inhibitory neurons), followed 7 weeks later by injection of EnvA-pseudotyped RabΔG-H2B-EGFP into contralateral VPL thalamus (3 mice). As seen with the first approach, monosynaptic cortical inputs arise almost exclusively from contralateral primary (SSp; ii) and supplemental (SSs; iii) somatosensory cortices, but not primary motor cortex (MOp) or rostral sensorimotor cortex (rSM; i). Serial two-photon tomography was used to quantify other monosynaptic inputs throughout the brain (see Materials and Methods and **Suppl. Table 1**). **(B)** Monosynaptic retrograde rabies tracing from Cu inhibitory neurons through injection of AAV-DIO-TVA-G into the Cu region of *VGAT-Cre* mice, followed 3-4 weeks later by injection of EnvA-pseudotyped RabΔG-H2B-EGFP into the Cu region (3 mice). Monosynaptic cortical inputs also arise from contralateral SSp (v) and SSs (vi), but also include a large population of corticofugal neurons throughout contralateral rSM (iv). Serial two-photon tomography was used to quantify other monosynaptic inputs throughout the brain (see Materials and Methods and **Suppl. Table 2**). **(C)** Control experiment for disynaptic retrograde rabies tracing in **Fig. 5C**. For monosynaptic transport from CL neurons, AAVretro-FlpO was injected into the contralateral VPL thalamus and Flp-dependent rabies helper viruses AAV-fDIO-TVA and AAV-fDIO-G were injected into the Cu region. After 3-4 weeks, EnvA-pseudotyped RabΔG-H2B-EGFP (expressing nuclear-localized EGFP) was injected into contralateral VPL thalamus, selectively targeting CL neurons that had received helper viruses. To prevent disynaptic transport, supplemental G-protein was not provided to local Cu inhibitory neurons. Mirroring the results of monosynaptic tracing from CL neurons (**Fig. 5A**, left), cortical inputs arise almost exclusively from contralateral SSp and SSs, with little to no labeling present in rSM. As expected, labeling was also found in cervical DRG and in presumptive local Cu inhibitory neurons (not shown). (4 mice).

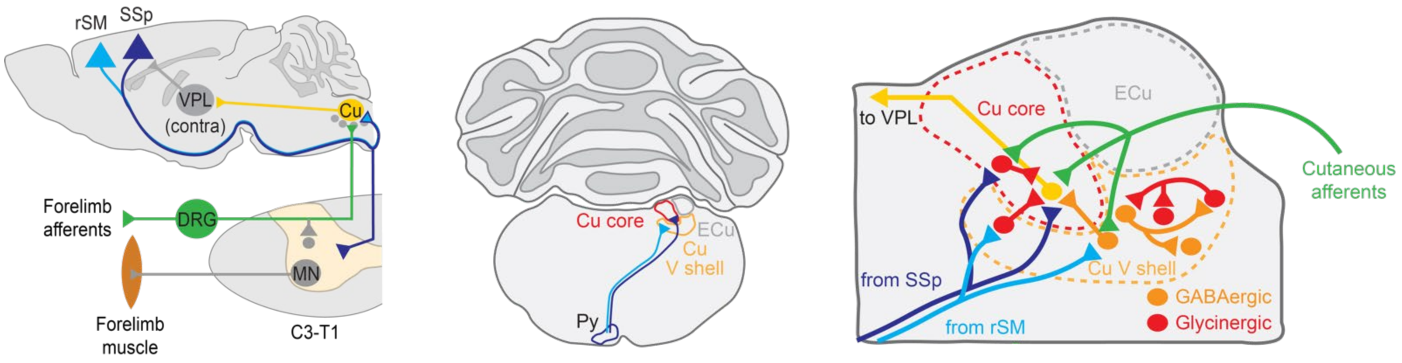


**Supplementary Figure 10. Summary of local and long-distance cuneate circuits.**
The main cuneate nucleus is a major conduit of forelimb sensory information to supraspinal regions, including the neocortex, via cuneolemniscal projections (yellow) to the thalamus (VPL). The core region of the middle cuneate (Cu) receives direct input from cutaneous afferents (green) that innervate the glabrous pad of the hand and reside in the dorsal root ganglia (DRG). GABAergic neurons (orange) located largely in the cuneate ventral shell (V shell) and glycinergic neurons (red) located in the cuneate ventral shell and cuneate core directly inhibit cuneolemniscal neurons. These inhibitory neurons receive inhibitory input, potentially from local connections, and also receive direct input from ascending cutaneous afferents. The cuneate core region is heavily targeted by corticospinal neurons residing in primary somatosensory cortex (SSp; dark blue), and cuneolemniscal neurons that reside in this core region receive direct input from SSp projections. Inhibitory cuneate neurons also receive SSp input, but unlike cuneolemniscal neurons, are also targeted by corticofugal neurons in rostral sensorimotor cortex (rSM; light blue), which do not project to the cuneate core, but rather target the shell region ventral to the cuneate. Cuneate inhibitory circuits provide a means for bidirectional modulation of the transmission of tactile information through the cuneate core, and their activation or inhibition perturbs the execution of tactile-guided dexterous behaviors. ECu, external cuneate; Py, pyramidal tract.

**Supplementary Movie 1. Normal execution of the string pull task.**
Cu inhibitory neurons were targeted for optogenetic activation by unilateral injection of AAV-DIO-oChIEF-Citrine into the Cu region of *VGAT-Cre* mice followed by implantation of an optical fiber. With the light off, behavioral performance is normal as both hands exhibit smooth cycles with uninterrupted pulling paths. Automated markerless tracking of the hands (Mathis, Mamidanna et al. 2018, Nath, Mathis et al. 2019) was used for kinematic quantification (see **Fig. 3A-D**).

**Supplementary Movie 2. Activation of cuneate inhibitory circuits disrupts string pull performance.**
In the same animal shown in **Supplementary Movie 1**, photostimulation (473 nm, 10 Hz, 50 msec pulse width) results in frequent ipsilateral prehension mistakes (left hand) and multiple grasp attempts at the top of the string pull cycle (see **Fig. 3A-D**).

**Supplementary Movie 3. Suppression of cuneate inhibitory circuits can cause premature termination of string pull behavior.**
Cu inhibitory neurons were targeted for optogenetic inhibition by injecting AAV-SIO-stGtACR2-FusionRed into the Cu region of *VGAT-Cre* mice followed by implantation of an optical fiber. In a subset of mice, photoinhibition (473 nm, continuous) causes early termination of a pulling bout and clutching of the ipsilateral (right) hand.

**Supplementary Table 1. Brain-wide monosynaptic inputs to cuneolemniscal neurons.**
Serial two-photon tomography was used to quantify monosynaptic inputs to cuneolemniscal neurons throughout the brain (see **Suppl. Fig. 9A** and Materials and Methods). Ipsilateral and contralateral quantification (relative to the injection site) is displayed in separate tabs for data collected from 3 mice. Columns display: region volumes; mean raw intensity of fluorescence in each region across mice; mean intensity of fluorescence per cubic mm in each region; SEM of mean intensity of fluorescence per cubic mm in each region; the mean probability normalized to the highest whole-brain mean intensity per cubic mm; and the SEM of the mean probability normalized to the highest whole-brain mean intensity per cubic mm. All data can be sorted by column values. Region abbreviations can be found at https://mouse.brain-map.org/static/atlas.

**Supplementary Table 2. Brain-wide monosynaptic inputs to cuneate inhibitory neurons.**
Serial two-photon tomography was used to quantify monosynaptic inputs to Cu inhibitory neurons throughout the brain (see **Suppl. Fig. 9B** and Materials and Methods). Ipsilateral and contralateral quantification (relative to the injection site) is displayed in separate tabs for data collected from 3 mice. Columns display: region volumes; mean raw intensity of fluorescence in each region across mice; mean intensity of fluorescence per cubic mm in each region; SEM of mean intensity of fluorescence per cubic mm in each region; the mean probability normalized to the highest whole-brain mean intensity per cubic mm; and the SEM of the mean probability normalized to the highest whole-brain mean intensity per cubic mm. All data can be sorted by column values. Region abbreviations can be found at https://mouse.brain-map.org/static/atlas.
